# A TBP-independent mechanism for RNA Polymerase II transcription

**DOI:** 10.1101/2021.03.28.437425

**Authors:** James Z.J. Kwan, Thomas F. Nguyen, Marek A. Budzyński, Jieying Cui, Rachel M. Price, Sheila S. Teves

## Abstract

Transcription by RNA Polymerase II (Pol II) is initiated by the hierarchical assembly of the Pre-Initiation Complex onto promoter DNA. Decades of *in vitro* and yeast research have shown that the TATA-box binding protein (TBP) is essential to Pol II initiation by triggering the binding of other general transcription factors, and ensuring proper Pol II loading. Here, we report instead that acute depletion of TBP in mouse embryonic stem cells (mESCs) has no global effect on ongoing Pol II transcription. Surprisingly, Pol II transcriptional induction through the Heat Shock Response or cellular differentiation also occurs normally in the absence of TBP. In contrast, acute TBP depletion severely impairs initiation by RNA Polymerase III. Lastly, we show that a metazoan-specific paralog of TBP is expressed in mESCs and that it binds to promoter regions of active Pol II genes even in the absence of TBP. Taken together, our findings reveal an unexplored TBP-independent process in mESCs that points to a diversity in Pol II transcription initiation mechanisms.

## Introduction

Transcription is the first step in gene expression, and it begins with the stepwise assembly of general transcription factors (GTFs) and RNA Polymerase II (Pol II) to form the Pre-Initiation Complex (PIC) at the promoters of genes (1). First, TATA-box binding protein (TBP), a core member of the GTF TFIID, binds onto gene promoters (1, 2). The conformational changes induced by TBP and TFIID trigger the binding of TFIIA and TFIIB (3), followed by TFIIF and Pol II, which further stabilizes the partial PIC (3). Lastly, the binding of TFIIE and TFIIH completes the PIC, where the enzymatic activity of TFIIH unwinds DNA to form a transcription bubble, allowing for Pol II progression and the incorporation of incoming nucleotides into the nascent RNA (4).

Decades of research using yeast genetics combined with *in vitro* reconstitution of yeast and human Pol II complexes have revealed a central role for TBP in the transcription initiation process. First identified as part of TFIID, TBP recognizes and binds to the TATA sequence in gene promoters (5, 6). The saddle-shaped protein forms extensive contact with promoter DNA, resulting in a 90° bend and partial unwinding of the double helix (3, 7, 8). TBP is highly conserved throughout Eukarya (8, 9). The C-terminal DNA binding domain of TBP consisting of 181 amino acids can be found from Archaea to Humans (10–12). Indeed, TBP purified from *Saccharomyces cerevisiae* can substitute for mammalian TFIID in *in vitro* transcription systems (13, 14).

Early studies in *S. cerevisiae* expressing TBP mutations show that the loss of functional TBP led to decreased GTF and Pol II recruitment and transcription *in vivo* (2, 15, 16). More recently, acute TBP depletion in yeast cells via the anchor-away system led to a rapid, genome-wide loss of Pol II and GTF occupancy (17, 18). In addition to facilitating Pol II transcription, TBP is also involved in at least two other RNA polymerases found in eukaryotes, and conditional inactivation of TBP in yeast led to a dramatic decrease in both RNA Pol I and Pol III transcription activity (2, 19–22).

Its crucial role in transcription combined with its high conservation have led to the model that TBP is universally essential for transcription (23). However, studies of mammalian TBP led to findings that were inconsistent with this hypothesis. While TBP deletion in mice is embryonic lethal, mouse embryos harbouring a homozygous TBP knock out were viable up to ~40-cell stage of embryogenesis and still displayed high levels of Pol II transcription (24). Furthermore, acute depletion of TBP in mouse embryonic stem cells (mESCs) showed that global transcription levels remained unaffected (25). These studies hint at an undefined role for TBP in mammalian cells that differ from its well-defined role in yeast cells.

A potential answer may lie in the emergence of TBP paralogs with specialized cell-type functions throughout evolution. In some species, these paralogs have been found to bind onto genomic loci distinct from TBP and to regulate cell-type specific genes (26, 27). For example, TBP-related factors TRF1 and TRF2 were first discovered in *Drosophila* with TRF1 being able to substitute for TBP in TATA transcriptional systems (28–31). A third vertebrate-specific paralog, TRF3, was found from fish to humans, but absent from *Drosophila and C. elegans* (32). TRF3 was also recently reported to mediate transcription of oocyte-specific genes and can form a non-canonical complex with TFIIA/TFIIB in mouse oocytes (33).

In this study, we re-examined the role of TBP in transcription initiation in mESCs. We report that, inconsistent with a requirement for TBP in Pol II-dependent transcription, rapid depletion of TBP has no global effect on nascent RNA levels and Pol II occupancy on chromatin. In contrast, tRNA transcription and Pol III chromatin occupancy are severely impacted upon loss of TBP. We found that transcription induction via heat shock and retinoic acid-mediated differentiation is retained in the absence of TBP. Lastly, we show that TRF2 interacts with Pol II and shares binding sites with TBP, and that, upon depletion of TBP, TRF2 remains bound to these genomic regions. These results provide evidence for a TBP-independent Pol II transcription in mESCs.

## Results

### TBP depletion does not affect global Pol II Transcription

We have previously generated a homozygous knock-in of the minimal auxin-inducible degron at the TBP locus (mAID-TBP) in mESCs (C64 cells) (34), which enables rapid and acute depletion upon the addition of indole-3-acetic acid (IAA) (Figure 1A) (35, 36). Treatment of C64 cells with 6 hours of IAA results in a ~90% depletion of mAID-TBP, as measured by Western blot and immunofluorescence (IF) analyses using an α-TBP antibody, with no significant effects on cell viability (Figure 1B, Supplemental Figure 1A-C). To assess the acute depletion of TBP genome-wide, we performed Cleavage Under Targets and Tagmentation (CUT&Tag) analysis for TBP (37). In CUT&Tag, antibodies to the protein of interest recruits the Protein A (pA)-Tn5 fusion to specific loci, enabling for efficient chromatin profiling of DNA binding proteins. After spike-in normalization, CUT&Tag analysis of TBP under control conditions shows high enrichment of TBP at transcription start sites (TSSs) (Figure 1C-D), with high reproducibility across replicates (Supplemental Figure 1D-E), consistent with its role in promoter recognition and binding. After 6 hours of IAA treatment, we observed a near complete depletion of bound TBP on promoters of two housekeeping genes, *Gapdh* and *Actb* (Figure 1C, Supplemental Figure 1D). To assess the magnitude of TBP depletion genome-wide, first we displayed the CUT&Tag signal in a 2 kb region surrounding the TSS for all genes as heatmaps, with genes arranged by decreasing Pol II signal in control conditions, and as average plots for all TSS (Figure 1D, Supplemental Figure 1E). We also quantified the read counts for TBP occupancy in a 1 kb region surrounding the TSS of all genes with and without IAA treatment and displayed as a scatter plot (Figure 1E). For both, we observed over 90% decrease in CUT&Tag signal at TSSs after IAA treatment genome-wide.

**Figure 1.**
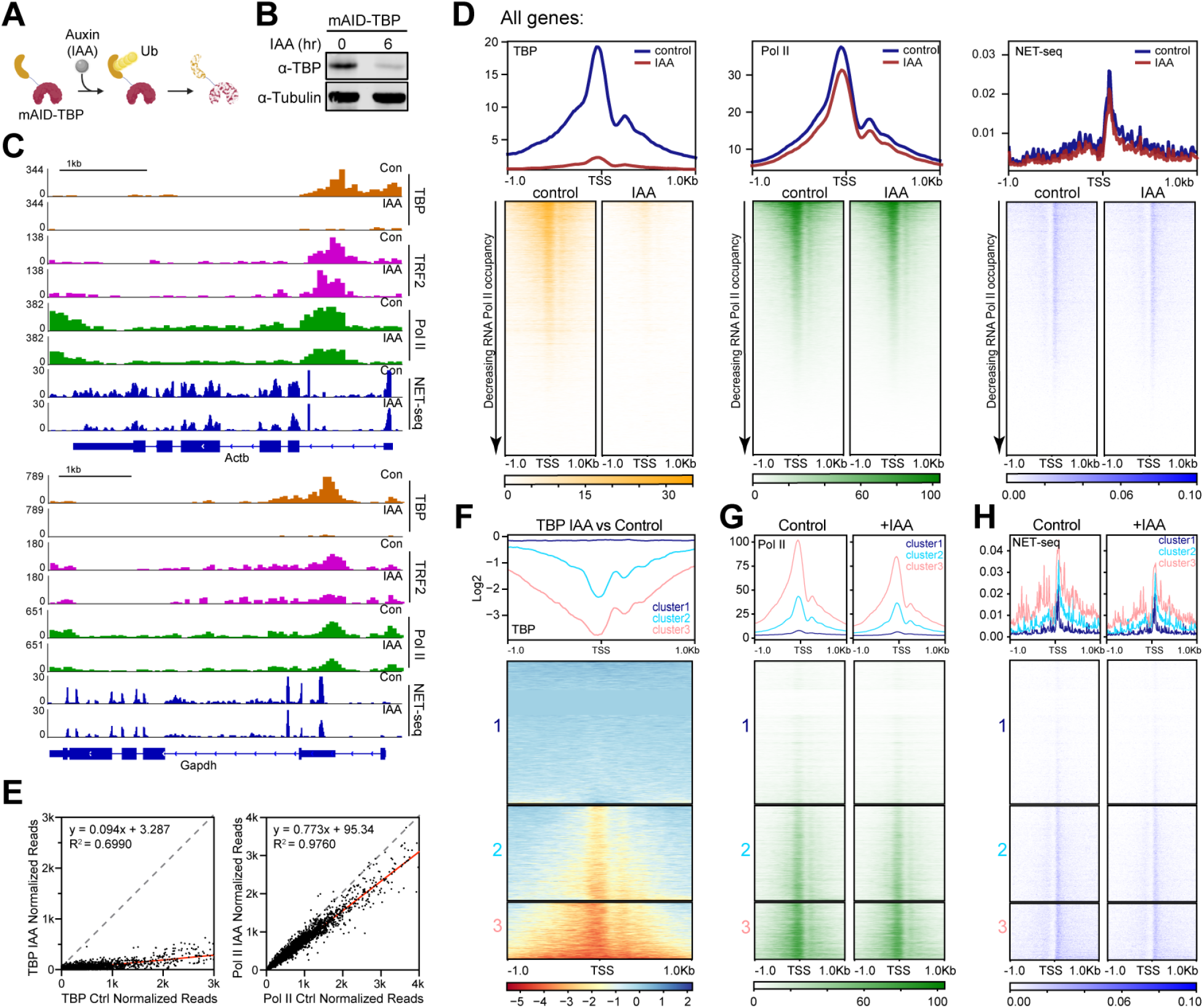
Global Pol II-mediated transcription is TBP-independent in mouse embryonic stem cells. **(A)** Cartoon of auxin-inducible degradation of TBP. mAID is endogenously knocked into TBP through CRISPR-Cas9 gene editing. **(B)** Western blot analysis of auxin (IAA)-mediated degradation of TBP in mESCs with the homozygous knock-in of mAID in the *TBP* locus (C64 mESCs). For all IAA-treated C64 mESCs, cells were incubated with 500 μM IAA for 6 hours unless otherwise stated. Control samples were treated with DMSO for 6 hours **(C)** Gene browser tracks at *Actb* (top) and *Gapdh* (bottom) for CUT&Tag analyses of TBP (orange), TRF2 (magenta) and Pol II (green), and strand-specific reads from NET-seq data (blue) in control (Con) or IAA-treated (IAA) C64 mESCs. **(D)** Genome-wide average plots (top) and heatmaps arranged by decreasing Pol II occupancy (bottom) of TBP CUT&Tag (left), Pol II CUT&Tag (middle), and NET-seq (right) on a 2kb window surrounding the TSS of all genes for control and IAA treated C64 mESCs. **(E)** Read counts of TBP control vs IAA-treated C64 mESCs were summed for the 1kb window surrounding the TSS of each gene, displayed as a scatter plot (left). Similar analysis for Pol II control vs IAA-treated samples (right). **(F)** k-means clustering with k=3 of the log_2_ IAA/Control of TBP CUT&Tag data. **(G-H)** Pol II CUT&Tag (G) and NET-seq (H) average plots and heatmaps by k=3 clustering from (F).

To investigate the precise role of TBP in mESCs, we examined the effects of TBP depletion on genome-wide Pol II occupancy in C64 cells using CUT&Tag. After spike-in normalization, we observed that, in control samples, Pol II binds at high levels on promoters and gene bodies of *Gapdh and Actb* (Figure 1C, Supplemental Figure 1D). Surprisingly, after 6 hours of IAA treatment (+ IAA), Pol II occupancy at the *Gapdh* and *Actb* loci remained unchanged. We then plotted the genome-wide occupancy of Pol II surrounding the TSS for all genes as heatmaps, with genes arranged by decreasing Pol II levels (Figure 1D, Supplemental Figure 1E). We also displayed the read counts for Pol II occupancy on all gene bodies with and without IAA treatment as a scatter plot (Figure 1E). As expected, control conditions show Pol II binding for active genes. Strikingly, we observed largely no global change in genome-wide Pol II occupancy after TBP depletion (Figure 1D-E, Supplemental Figure 1E), inconsistent with a requirement for TBP in Pol II recruitment to promoters.

To directly measure Pol II activity, we performed NET-seq (native elongating transcript sequencing) analysis on control and TBP-depleted mESCs. NET-seq allows for the capture and mapping at single nucleotide resolution of newly transcribed RNA from elongating Pol II (38) through an extensive cellular fractionation method that separates chromatin-bound Pol II from the cytoplasm and nucleoplasm. After normalization with spike-in controls, we detected comparable intronic and exonic signals in *Gapdh* and *Actb* genes for both control and IAA-treated samples (Figure 1C, Supplemental Figure 1D). We then displayed normalized NET-seq reads as heatmaps in a 2 kb region surrounding the TSS for all genes (Figure 1D, Supplemental Figure 1E). Globally, we observe no distinguishable differences in nascent RNA levels between control and IAA-treated cells. Consistent with what was observed for Pol II occupancy, our NET-seq data show that acute TBP depletion does not lead to the disruption of genome-wide Pol II-mediated transcription in mESCs.

We next calculated the log-fold change in TBP signal in IAA-treated samples relative to control, displayed the values as a heatmap, and performed unbiased k-means clustering with k=3 (Figure 1F). Cluster 3 showed the most significant amount of TBP depletion while Cluster 1 showed no changes in TBP depletion upon IAA treatment. We then plotted the Pol II CUT&Tag and NET-seq data for these three clusters (Figure 1G-H). Pol II occupancy and nascent RNA levels for Cluster 3 showed no changes in IAA condition compared to control, indicating that although these genes experience the largest decrease in TBP occupancy, transcriptional activity is not perturbed. Taken together, these results show that Pol II transcription in mESCs is largely TBP independent.

### TBP is dispensable for transcription of specific classes of genes, Pol II pausing, and enhancer RNA synthesis

Our results showing that Pol II transcription is independent of TBP in mESCs are in stark contrast with the well characterized function of TBP in budding yeast cells. We hypothesized that TBP might act preferentially to promote the transcription of mouse genes that share a large degree of homology with budding yeast genes. *S. cerevisiae* and *M. musculus* protein-coding genes were queried for homologous and non-homologous genes using MouseMine, an online resource that mines data from multiple model organism databases (39). From the TBP CUT&Tag data, we observed that TBP binds to both homologous (4475 genes) and non-homologous genes (15528 genes) equally well and that TBP depletion is near complete for both sets (Supplemental Figure 2D-E). We then displayed as scatter plots the normalized CUT&Tag read counts for Pol II versus TBP in control samples for homologous and non-homologous genes (Supplemental Figure 2A-B). We observed a positive relationship between Pol II and TBP occupancy in both subsets of genes (R^2^ = 0.4137 and 0.4902 respectively), which is similar to the relationship for all genes (Supplemental Figure 2C, R^2^ = 0.4723). This finding suggests that TBP occupancy and transcriptional activity are related regardless of whether the gene shares homologous with *S. cerevisiae*. Next, we plotted the genome-wide occupancy of Pol II in control and IAA treated cells surrounding the TSS for homologous (Figure 2A) and non-homologous (Supplemental Figure 2E) genes as heatmaps, with genes arranged by decreasing Pol II levels. We observed no changes in Pol II occupancy after TBP depletion for either subset of genes. To directly observe the activity of Pol II across these gene subsets, we plotted normalized NET-seq reads in control and IAA treated cells as heatmaps in a 2 kb region surrounding the TSS for yeast homologous genes (Figure 2A) and non-homologous (Supplemental Figure 2E) genes as heatmaps. Consistent with our Pol II CUT&Tag analyses, we observe no differences in nascent RNA levels between control and TBP depleted cells across yeast homologous and non-homologous genes. Our results indicate that TBP depletion does not affect gene transcriptional activity, regardless of the gene’s genomic divergence from *S. cerevisiae*.

**Figure 2.**
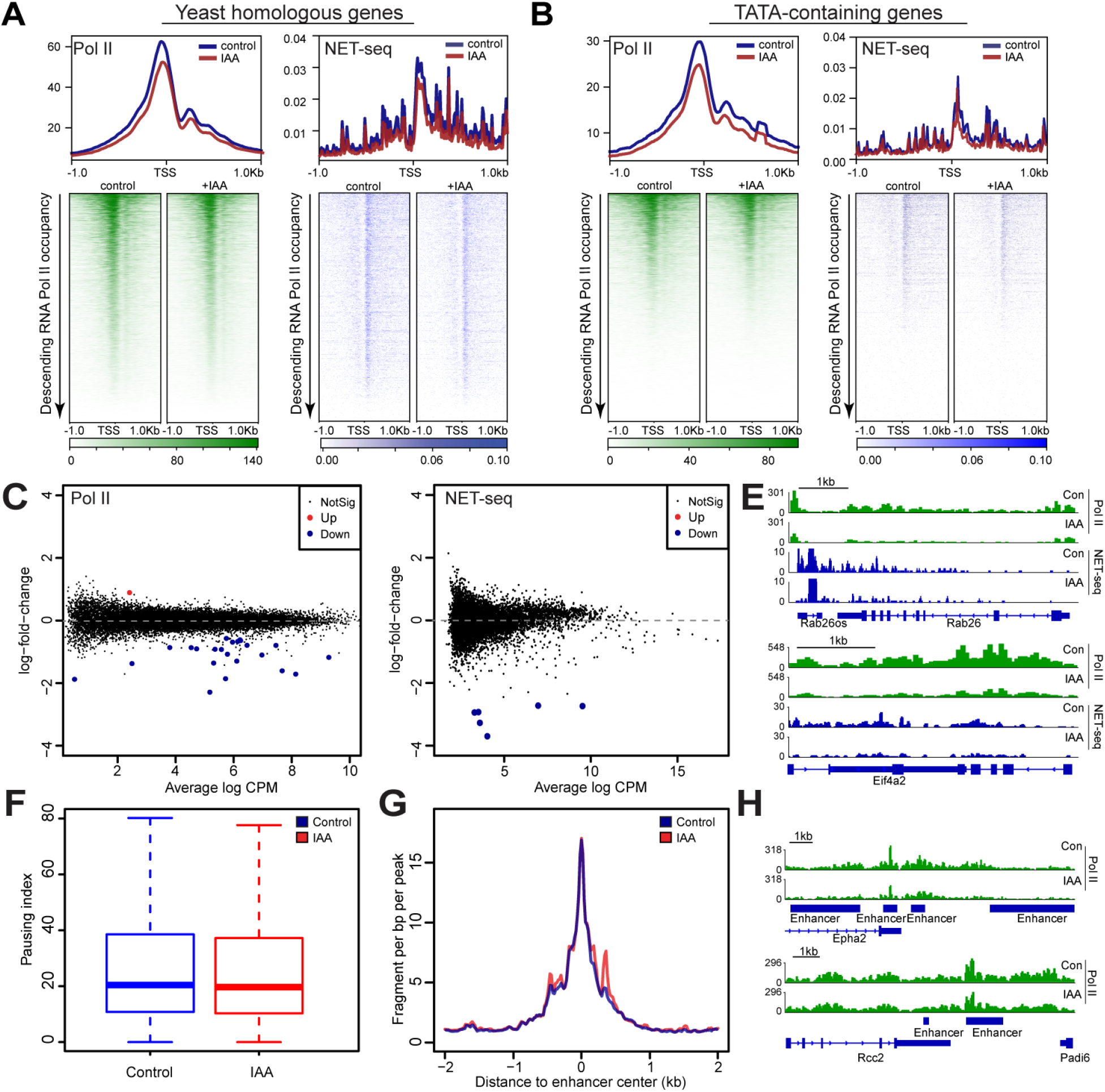
Transcription of yeast homologous and TATA-containing genes, Pol II pausing, and enhancer RNA synthesis is TBP-independent in mouse embryonic stem cells. **(A-B)** Average plots (top) and heatmaps arranged by decreasing Pol II occupancy (bottom) of Pol II CUT&Tag (left) and NET-seq (right) on a 2kb window surrounding the TSS of all yeast homologous genes (A) and TATA-containing genes (B) for control and IAA-treated (IAA) C64 mESCs. **(C-D)** Differential gene analysis of Pol II CUT&Tag (C) and NET-seq analysis (D) on all genes in control vs. IAA-treated C64 mESCs. **(E)** Gene browser tracks for *Rab26* and *Eif4a2* for Pol II CUT&Tag analyses (green) and strand specific reads from NET-seq data (blue) in control (Con) or IAA-treated C64 mESCs. **(F)** Average pausing index of Pol II across all genes in control (blue) and IAA treatments (red) as calculated from Pol II CUT&Tag data **(G)** Pol II CUT&Tag signals around the center of enhancers in control (blue) and IAA-treated (red) C64 mESCs. **(H)** Gene browser tracks for different enhancer regions for Pol II CUT&Tag analysis in control (Con) or IAA-treated C64 mESCs.

TBP was first identified to bind to the canonical TATA sequence (TATAWAW), which is present in about ~3-15% of promoters (7, 40–43). Despite its moniker, TBP is required for transcription of both TATA-containing and TATA-less promoters in yeast cells, though its role in TATA-less yeast promoters remains more nuanced (41, 44). We hypothesized that mammalian TBP may act preferentially to promote the transcription of either TATA-containing or TATA-less promoters. We therefore examined the effects of TBP depletion in these two promoter classes. We grouped all annotated TSS/promoters as either TATA-containing (5357 genes) or TATA-less (28312 genes), and plotted the TBP (Supplemental Figure 2F-G) and Pol II (Figure 2B) CUT&Tag data as normalized heatmaps and average plots before and after TBP depletion in a 2kb region surrounding the TSS. In control cells, the average Pol II occupancy levels are roughly similar for both TATA-containing and TATA-less genes (Figure 2B, Supplemental Figure 2G). Similarly, TBP binds to promoters of both gene sets roughly equally (Supplemental Figure 2F-G). Upon TBP depletion, we observed a similar decrease in bound TBP in both gene sets (Supplemental Figure 2F-G), whereas Pol II occupancy remained unchanged in both promoter classes (Figure 2B, Supplemental Figure 2G, Supplemental Figure 3A). We performed a similar analysis for our NET-seq data and observed no global change of transcription activity for both TATA-containing (Figure 2B, Supplemental Figure 3A) and TATA-less genes (Supplemental Figure 2G, Supplemental Figure 3A) upon TBP depletion. These results indicate that, regardless of the presence or absence of the TATA binding motif, Pol II activity is unaffected by TBP depletion.

Global analyses may mask effects on individual genes. To identify if individual genes are TBP-dependent, we used the edgeR Bioconductor package, which includes the differential gene analysis tool that can be used to identify genes with statistically significant differences (45) in Pol II occupancy (CUT&Tag, Figure 2C) or activity (NET-seq Figure 2D) between control and IAA-treated samples by comparing normalized and filtered read counts. Consistent with global average plots and heatmaps, differential gene analyses of both Pol II and NET-seq data in control vs IAA-treated samples show that 99.9% of genes do not display significant changes upon TBP depletion (black dots in Figure 2C-D). However, 23 genes showed significant downregulation with a ~2-4 fold decrease and 1 gene with a ~2 fold increase in Pol II binding activity upon IAA treatment, with a mix of protein-coding and long non-coding RNA genes (Supplemental Figure 3B). Specifically, gene browser tracks for Pol II occupancy and NET-seq signal on *Rab26* and *Eif4a2* genes show high levels under control conditions, and decreased levels for both Pol II occupancy and NET-seq signal after IAA treatment, suggesting that these genes are either directly or indirectly affected by TBP depletion (Figure 2E, Supplemental Figure 3C). In contrast to the global effects of TBP depletion in yeast cells, these findings show that only a select number of genes are affected upon TBP depletion in mouse ESCs.

Promoter-proximal pausing of Pol II is an important transcriptional regulatory step in many active genes. As Pol II escapes the PIC complex and begins transcribing, it pausest 40-60 nucleotides downstream of the TSS and stalls for further signal (46–49). To investigate whether promoter-proximal pausing is affected upon loss of TBP, we measured the “Pausing Index”, also known as “Traveling Ratio”, from our Pol II CUT&Tag data (Figure 2F). The Pausing Index (PI) is defined as the ratio of Pol II accumulation at the promoter relative to Pol II levels within gene bodies and was calculated using the NRSA suite (50–53). The average PI of each gene was calculated and the median values were then taken for control and IAA treated samples which showed similar values. A pausing index of 20.4 for control vs 19.7 for IAA-treated samples (Figure 2F), with individual replicates displaying very similar results (Supplemental Figure 3D), indicates that Pol II pausing is also not affected by depletion of TBP.

Pol II-transcription also occurs at enhancers, resulting in the production of enhancer RNAs (eRNAs) (54, 55). To investigate whether TBP depletion affects eRNAs we performed enhancer analysis using the NRSA suite, which also has the ability to search for *de novo* enhancers through detection of bi-directional non-coding transcription, a direct and reliable indicator of enhancer activity (53). When we plotted the Pol II CUT&Tag signal on these *de novo* identified enhancers, we observed no change in Pol II signal for control and IAA treated samples with individual replicates displaying similar results, indicating that transcription at enhancers are also TBP-independent (Figure 2G-H, Supplemental Figure 3E).

### TBP depletion does not affect gene activation

We have shown that acute depletion of TBP has no global effect on Pol II transcription of active genes. We next examined if TBP is required for the transcriptional activation of silent genes. To address this question, we used a classical transcription induction system in eukaryotic cells: the heat shock (HS) response. A highly conserved protective mechanism to stressors like elevated temperatures, reactive oxygen species, and heavy metals, the HS response includes the rapid transcription induction of HS protein genes (56–60), which act as molecular chaperones to facilitate protein refolding or degradation resulting from stress (61–64). Transcription of HS genes is regulated at the level of Pol II pausing (65, 66). Upon HS, the paused Pol II is then released into active elongation (67–69). To induce the transcription of HS genes, we exposed C64 cells to heat shock at 42°C for 30 minutes (Figure 3A) and performed Pol II CUT&Tag and NET-seq analyses. Gene browser tracks for CUT&Tag and NET-seq at classical heat shock genes *Hspa1a, Hsp90aa1, Dnajb1*, and *Rbm39* gene loci show massive changes in Pol II binding and activity upon HS compared to control samples with a 3-4 fold increase for upregulated genes and 2 fold decrease for the downregulated gene. (Figure 3B, Supplemental Figure 4D-E). Differential gene analysis of Pol II CUT&Tag data shows 577 and 503 genes are significantly upregulated and downregulated, respectively, in HS samples compared to control (Figure 3C). The significantly upregulated genes from differential gene analysis include several well-known HS genes such as *Hsph1, Dnajb1* and *Hsp90aa1* (Supplemental Figure 4A). Gene ontology analysis shows that upregulated genes are enriched for unfolded protein binding, chaperone binding, and heat shock protein binding, which are typical markers for the HS response (Supplemental Figure 4B). Furthermore, we observed that the pausing index decreased in the heat shock samples compared to control when looking at the top upregulated HS genes, suggesting there is more Pol II occupancy in the gene body compared to the proximal promoter site (Supplemental Figure 4C). These analyses validated our methods used to detect HS-induced gene expression.

**Figure 3.**
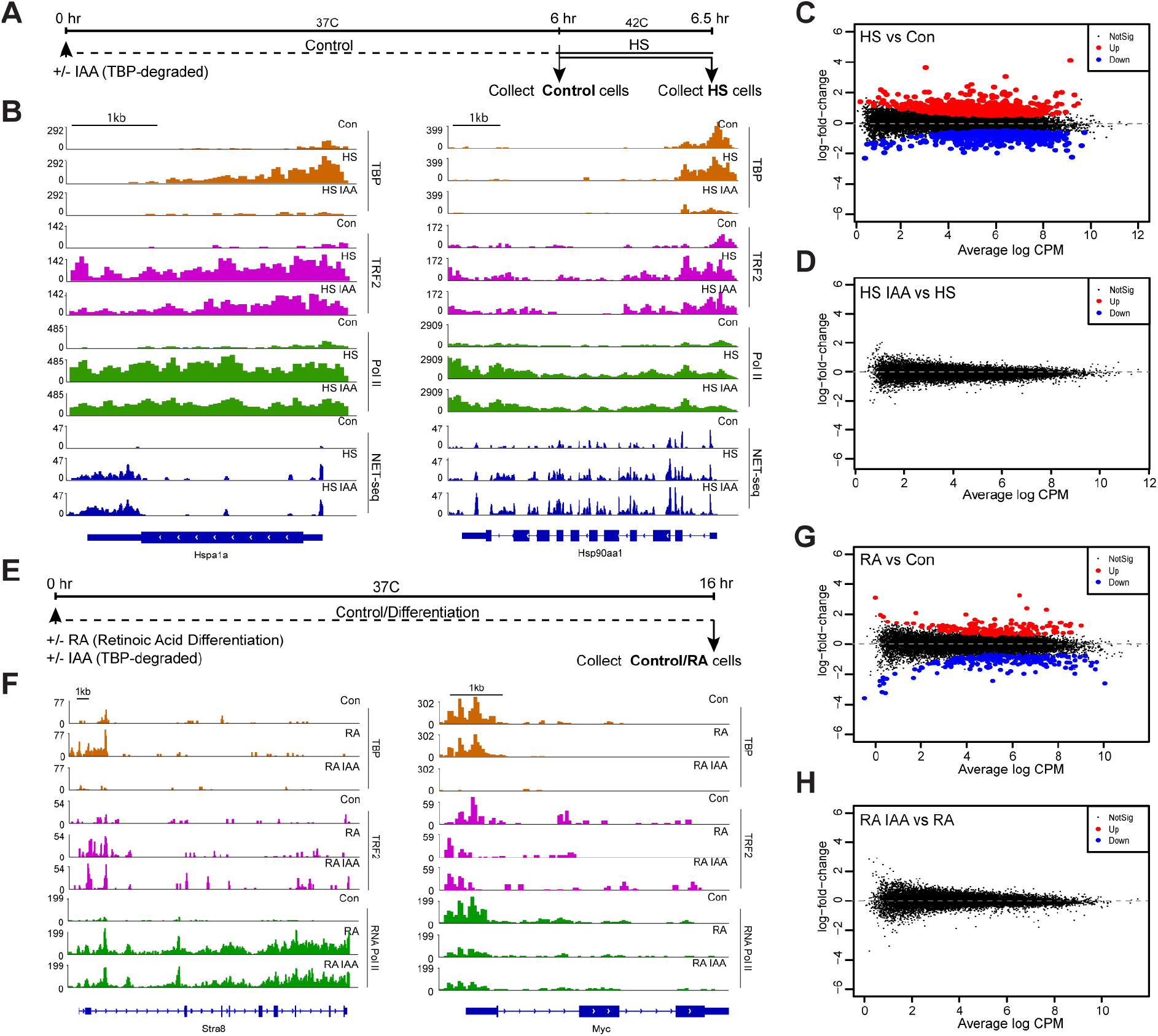
TBP is dispensable for gene activation by heat shock and retinoic acid differentiation in mESCs. **(A)** Schematic of heat shock treatments of mESCs. After 6 hours of treatment with auxin (IAA) or DMSO, cells were either collected or heat shocked at 42°C for 30 minutes before collection. **(B)** Gene browser tracks for *Hspa1a* (left) and *Hsp90aa1* (right) for CUT&Tag analyses of TBP (orange), TRF2 (magenta) and Pol II (green), and strand specific reads from NET-seq data (blue) in control (Con) or IAA-treated C64 mESCs. **(C-D)** Differential gene analysis of Pol II occupancy on all genes in heat shocked vs. control C64 mESCs (C) and in heat shocked and IAA-treated vs. heat shocked C64 mESCs (D). **(E)** Schematic of retinoic acid treatment of mESCs. Cells were treated for 16 hours with DMSO or IAA and retinoic acid (RA) before collection. **(F)** Gene browser tracks for *Stra8* (left) and *Myc* (right) for CUT&Tag analyses of TBP (orange), TRF2 (magenta) and Pol II (green) in control (Con) or IAA-treated C64 mESCs. **(G-H)** Differential gene analysis of Pol II occupancy on all genes in RA-treated vs. control C64 mESCs (G) and in RA and IAA-treated vs. RA-treated C64 mESCs (H).

To investigate the effects of TBP depletion on the induction of HS genes, we treated C64 cells with IAA for 6 hours followed by heat shock for 30 minutes, and performed Pol II CUT&Tag and NET-seq analysis (Figure 3A). We confirmed TBP depletion following IAA treatment on HS genes using CUT&Tag (Supplemental Figure 4F-G). Then, we visualized Pol II CUT&Tag in *Hspa1a and Hsp90aa1* (Figure 3B) and *Dnajb1* (Supplemental Figure 4E) under control and IAA-treated samples and confirmed that Pol II is paused at the promoter regions under normal conditions and that the pausing is unaffected by TBP depletion. Under HS + IAA conditions, gene browser tracks for Pol II CUT&Tag and NET-seq at the *Hspa1a, Hsp90aa1* and *Dnajb1* gene loci show that these genes are still actively transcribed (Figure 3B, Supplemental Figure 4E). Differential gene analysis of Pol II CUT&Tag for HS versus HS + IAA samples show no significant changes in gene expression (Figure 3D), indicating that global changes in expression induced by HS occur in the absence of TBP. We also plotted Pol II occupancy for the top upregulated and downregulated genes identified from differential gene analysis (Figure 3C) as heat maps centered at the TSS (Supplemental Figure 4H-I). Surprisingly, Pol II levels do not change between HS and HS + IAA conditions for both of these gene sets. These results suggest that depletion of TBP does not impair Pol II pausing and release during the induction of HS genes.

Next, we examined the role of TBP in gene activation through a different method of gene induction, namely Retinoic Acid (RA)-mediated differentiation of mESCs. RA is a vitamin A metabolite that is essential for cell differentiation in embryonic development (70–73). Treatment of mESCs with RA induces cellular differentiation into neuroectoderm and extraembryonic endoderm-like cells, which leads to the silencing of pluripotency markers and activation of ectoderm-specific genes within 6-24h of RA treatment (70, 74). To measure gene activation via RA differentiation, we treated C64 cells with RA for 16 hours (Figure 3E), and performed Pol II CUT&Tag. Gene browser tracks of *Stra8* and *Cdx1* genes, known to be upregulated in RA treatment (75, 76), shows low levels of Pol II in the control sample, followed by significant increase in Pol II occupancy after RA treatment (Figure 3F, Supplemental Figure 5C). Additionally, the pluripotency marker *Myc* and a downregulated gene *Rif1*, which are expressed highly under control conditions, display a large decrease in Pol II occupancy under RA conditions (Figure 3F, Supplemental Figure 5D). To measure global changes, we performed differential gene analysis of Pol II CUT&Tag, which revealed a significant number of genes that change expression upon RA treatment, including 154 upregulated and 162 downregulated genes with several well known RA genes (Figure 3G, Supplemental Figure 5A). Furthermore, we observed a decrease in the pausing index of the top upregulated genes during RA treatment, suggesting an increase of Pol II occupancy compared to the proximal promoter (Supplemental Figure 5B).

We next asked whether TBP depletion would impair RA-mediated induction of genes by performing Pol II CUT&Tag in TBP depleted and RA-treated cells (Figure 3E). As before, we verified TBP depletion in this regimen using CUT&Tag analysis (Supplemental Figure 5E-F). Gene browser tracks for Pol II CUT&Tag at the *Stra8* gene show similar levels of Pol II binding in both RA + IAA and RA-only conditions, suggesting that these genes are induced similarly with and without TBP depletion (Figure 3F). Differential gene analysis of Pol II CUT&Tag data for RA and RA + IAA samples (Figure 3H) showed no significant difference between the two conditions on a gene-by-gene basis. Next, we plotted the Pol II occupancy for the significantly upregulated and downregulated genes extracted from differential gene analysis (Figure 3G) for RA and RA + IAA treatment as heat maps centered at the TSS (Supplemental Figure 5G-H). Both RA and RA + IAA conditions display similar levels of Pol II, confirming that depleting TBP does not affect either the silencing or, more importantly, the activation of RA-specific genes. Surprisingly, these results suggest that gene activation that occurs at the level of Pol II recruitment and initiation does not require TBP.

### TBP is required for RNA Pol III transcription of tRNA genes

Previous studies have shown that TBP is required for initiation of the three main eukaryotic RNA Polymerases (2, 21, 23, 77). Whereas Pol II transcribes protein coding genes, Pol III transcribes all of the tRNAs, the 5S ribosomal RNA, the U6 spliceosomal RNA and other groups of non-coding RNAs that are critical for cellular functions such as RNA and protein homeostasis and cell growth. Given our findings for Pol II, we next asked if TBP is required for Pol III transcription in mESCs. To address this question, we performed CUT&Tag analysis of Pol III with and without TBP degradation. As before, we validated TBP depletion in these samples (Figure 4A-C, Supplemental Figure6). In control cells, Pol III binds at high levels on tRNA genes *LeuCAA* and *GlnCTG*. After TBP depletion, we observed a drastic decrease in Pol III occupancy at these tRNA genes and across each individual replicate (Figure 4A, Supplemental Figure 8A). To examine the effects of TBP depletion on Pol III occupancy across all tRNAs, we plotted the genome occupancy of Pol III in a 2kb window surrounding the TSS of all tRNAs in control and TBP depleted cells. We observed a similar drastic decrease of Pol III occupancy across all tRNAs upon TBP depletion (Figure 4C, Supplemental Figure 8B). We also determined the read counts normalized as RPKM values for Pol III on tRNAs with and without IAA treatment (Figure 4B). If there are no changes, all points would lie near the diagonal. Instead, we observed that most points are well below the diagonal, indicating a major effect of TBP depletion on Pol III binding to tRNA genes.

**Figure 4.**
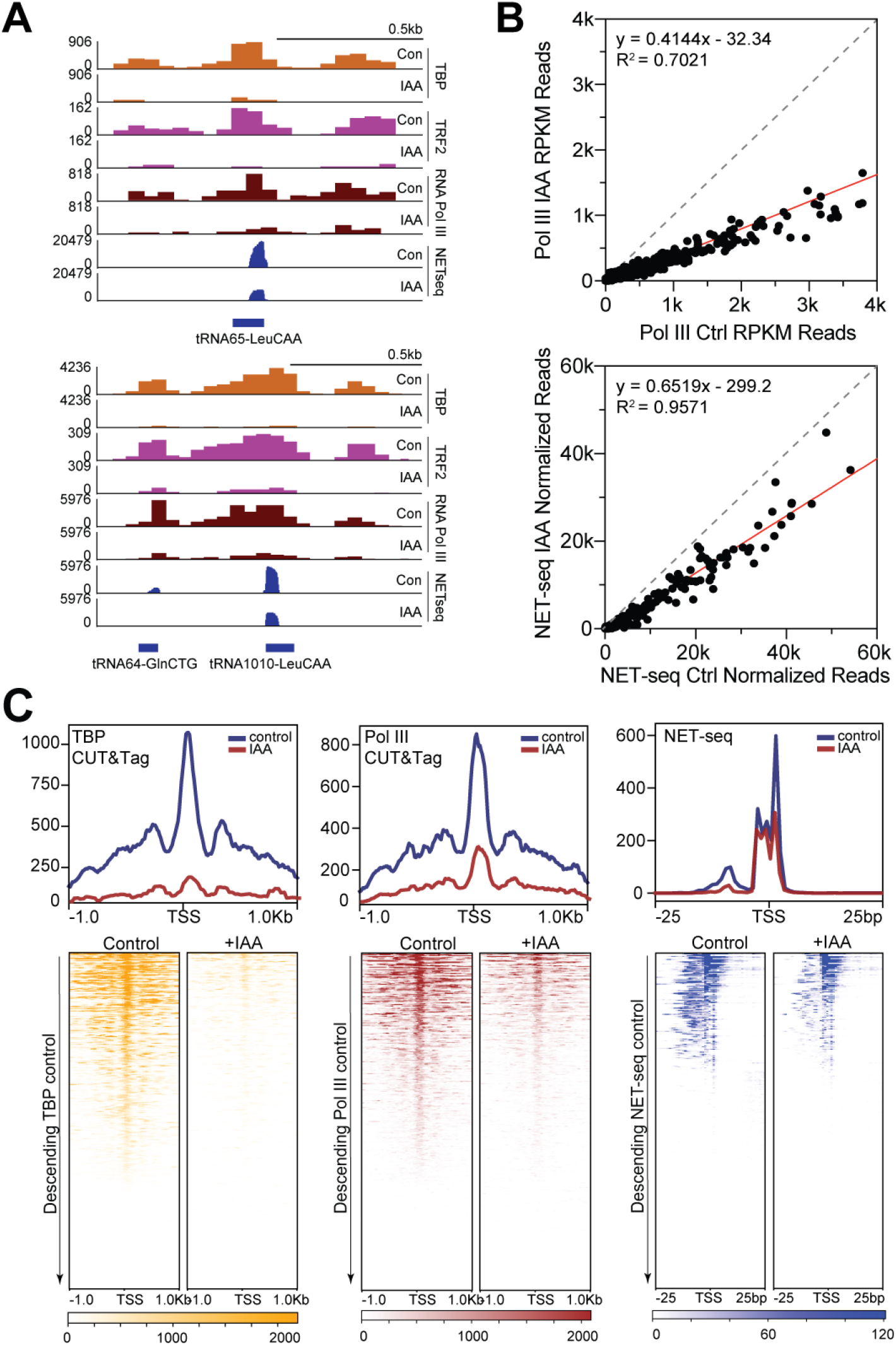
TBP is required for Pol III transcription of tRNAs in mouse embryonic stem cells. **(A)** Gene browser tracks at tRNA65-LeuCAA (top) and tRNA64-GlnCTG (bottom) for CUT&Tag analyses of TBP (orange), TRF2 (magenta), Pol III (maroon), and strand-specific reads from NET-seq data (blue) in control (Con) or IAA-treated (IAA) C64 mESCs. **(B)** Read counts in control vs. IAA-treated mESCs of Pol III occupancy across all tRNAs normalized by RPKM (top) and Pol III activity in a 50bp window surrounding the TSS of all tRNAs as measured by NET-seq (bottom). **(C)** Average plots (top) and heatmaps arranged by decreasing TBP occupancy (bottom) of TBP CUT&Tag (left), Pol III CUT&Tag (middle), and NET-seq (right) in a 2kb window (CUT&Tag) or 50 bp window (NET-seq) surrounding the TSS of all tRNAs.

As well as elongating Pol II, NET-seq can also capture the activity of elongating Pol III (78). To quantify Pol III activity, we analyzed NET-seq signals on tRNAs in control and TBP-depleted mESCs. After normalization with spike-in controls, we detected signals for tRNA genes *LeuCAA* and *GlnCTG* in control cells; however, upon IAA treatment, the signal at these tRNA genes is decreased 2-3 fold (Figure 4A, Supplemental Figure 8A). To examine the effects of TBP depletion across all tRNAs, we displayed normalized NET-seq reads as heatmaps in a 50 bp region surrounding the TSS for all tRNAs and observed a decrease of signal for all tRNAs upon TBP depletion (Figure 4C, Supplemental Figure 8B). We also determined the NET-seq read counts for all tRNAs with and without IAA treatment and displayed the data as a scatter plot (Figure 4B). Similar with the Pol III CUT&Tag analysis, the points mostly fall below the diagonal. The decrease in tRNAs upon TBP is more modest than the decrease of Pol III occupancy, which could be attributed to the abundance of mature tRNAs that may contaminate the NET-seq data. Nevertheless, what we observe in the NET-seq data is consistent with the change in Pol III occupancy on tRNA genes upon IAA treatment. Therefore, TBP is required for the transcription of tRNAs by Pol III in mESCs, consistent with previous studies in yeast and *in vitro* systems (22, 79, 80). Importantly, the consistent role of TBP in Pol III transcription contrasts strikingly with the TBP-independent Pol II transcription in mESCs.

### The TBP paralog TRF2 is expressed in mESCs and binds to promoters of active genes

The mouse genome contains two known TBP paralogs, *Tbpl1 and Tbpl2*, which code for the proteins TBP-related factor 2 and 3 (TRF2 and TRF3), respectively (11, 81, 82). We observed that TRF2 is expressed in mESCs, as confirmed through Western blot and IF analysis showing nuclear localization (Figure 5A Control Lane). Additionally, gene browser tracks for Pol II CUT&Tag at the genomic loci *Tbpl1* and *Tbpl2* indicate that *Tbpl1* is expressed at relatively high levels whereas *Tbpl2* falls within background signal (Supplemental Figure 7A). To test if TRF2 interacts with Pol II, we performed co-immunoprecipitation with pulldown for TRF2 and Western blot detection for Pol II. The presence of Pol II in the TRF2 pulldown suggests that these two proteins interact directly or indirectly (Figure 5B, left). We next examined whether TRF2 and TBP were interacting with each other. We performed co-immunoprecipitation with pulldown for TBP and Western blotted for TRF2 (Figure 5B, right). We detected no TRF2 in the TBP IP, suggesting these two factors do not interact with each other, and might not be found together in the same PIC.

**Figure 5.**
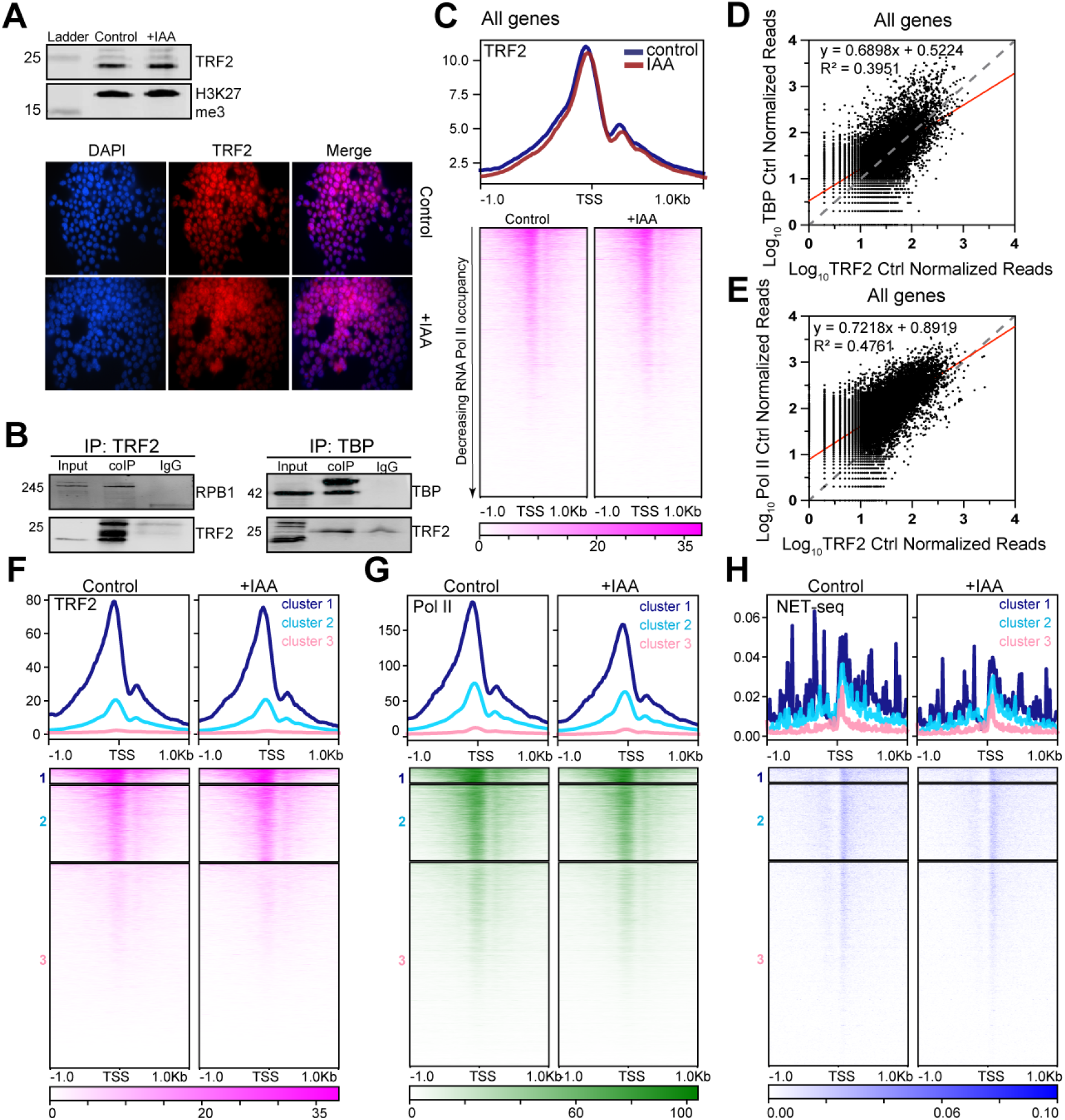
TBP paralog TRF2 is expressed in mESCs and binds to promoters of active genes. **(A)** Western blot analysis for α-TRF2 (21kDa) and α-H3K27me3 (17kDa) in control and IAA-treated C64 mESCs (top). Immunofluorescence imaging using α-TRF2 and DAPI staining (bottom) for control and IAA-treated conditions. **(B)** Co-Immunoprecipitation assay with pulldown using α-TRF2 antibody and blotting for RNA Pol II subunit RPB1 (left), or pulldown using α-TBP antibody and blotting for TRF2 (right). The TRF2 band is present in the left panel indicating interaction with RPB1 and absent in the right panel indicating a lack of interaction with TBP. **(C)** Genome-wide average plots (top) and heatmaps arranged by decreasing Pol II occupancy (bottom) of TRF2 CUT&Tag on a 2kb window surrounding the TSS of all genes for control and IAA treated C64 mESCs **(D-E)** Read counts of TRF2 control vs TBP control C64 mESCs were summed for the promoter of each gene (−250bp of TSS to TSS), log_10_ transformed and displayed as a scatter plot (top). Same analysis for TRF2 control vs Pol II control CUT&Tag data (bottom). Linear regression was done to determine slope and R^2^ value. **(F)** Genome-wide average plots (top) and heatmaps arranged by decreasing TRF2 occupancy (bottom) with k-means clustering with k=3 of control and IAA TRF2 CUT&Tag data. **(G-H)** Pol II CUT&Tag **(G)** and NET-seq **(H)** average plots and heatmaps by k=3 clustering from (F).

Where does TRF2 bind in the genome? To answer this question, we performed CUT&Tag for TRF2. At the *Actb* and *Gapdh* loci, we observed strong binding of TRF2 to promoters of these genes (Figure 1C, Supplemental Figure 7B). We then displayed the TRF2 CUT&Tag data as a heatmap and average plot for all genes, arranged by decreasing Pol II levels (Figure 5C, Supplemental Figure 7C). Similar to TBP, TRF2 binds to promoters of all active genes. Previous studies in various species have shown that TBP and its paralogs occupy and regulate distinct sets of genes (82–86). It was therefore surprising that in mESCs, TRF2 binds to all active genes, though the differences may reflect species-specificity. We further quantified the correlation between TBP and TRF2 by calculating the normalized log values for TBP and TRF2 CUT&Tag read counts at the promoters of all genes and displaying the values as a scatter plot (Figure 5D). Linear regression analysis shows a strong correlation between TRF2 and TBP occupancy (R^2^ = 0.3951), indicating that gene promoters that contain high levels of TBP also display high TRF2 occupancy. We performed a similar analysis between TRF2 and Pol II read counts (Figure 5E), and also observed moderate correlation (R^2^ = 0.4761). As with TBP, high levels of TRF2 correlates with high expression. Indeed, k-means clustering of TRF2 CUT&Tag reveals three clusters corresponding to high, medium, and low TRF2 occupancy (Figure 5F, Control). When Pol II CUT&Tag (Figure 5G, Control) and NET-seq data (Figure 5H, Control) are plotted according to the three TRF2 clusters, we confirmed that Pol II occupancy and activity correlates with TRF2 occupancy.

We next asked if TBP depletion affects TRF2 protein levels or its binding to promoters. We measured TRF2 protein levels by Western blot and IF with and without IAA treatment, and observed that global TRF2 levels remain unchanged upon IAA treatment (Figure 5A). Next, we performed TRF2 CUT&Tag on IAA-treated C64 cells to examine the genomic occupancy of TRF2 upon depletion of TBP. Gene browser tracks of *Actb* and *Gapdh* genes show that TRF2 binds to the promoter region in similar levels in the control and IAA-treated samples (Figure 1C, Supplemental Figure 7B). Furthermore, heatmaps for the global occupancy (Figure 5C, Supplemental Figure 7C) or the three clusters from k-means clustering (Figure 5F) of TRF2 in control and IAA-treated conditions show that TRF2 maintains promoter binding genome-wide upon TBP depletion. Importantly, Pol II CUT&Tag (Figure 5G) and NET-seq data (Figure 5H) show no change in IAA-treated samples compared to control in the three clusters of TRF2. Taken together, these results indicate that TRF2 binding is independent of TBP.

### TRF2 binds to all Pol II-promoter types and to tRNA genes in mESCs

Could TRF2 function redundantly with TBP in mESCs? To address this question, we examined the TRF2 occupancy on genes with the greatest change in TBP occupancy after depletion. Using the three clusters of log-fold change of TBP in IAA versus control samples, we plotted TRF2 CUT&Tag control and IAA data on clusters that show the greatest and least change in TBP occupancy levels upon IAA treatment (Figure 6A). This analysis shows that cluster 3, which includes genes with the greatest change in TBP levels upon IAA treatment, contains the highest level of TRF2, whereas cluster 1, which includes genes with the least change in TBP, shows the lowest TRF2 levels. Furthermore, TRF2 levels across the three clusters do not change upon IAA treatment. This analysis suggests that TRF2 could function redundantly with TBP for the majority of mouse genes.

**Figure 6.**
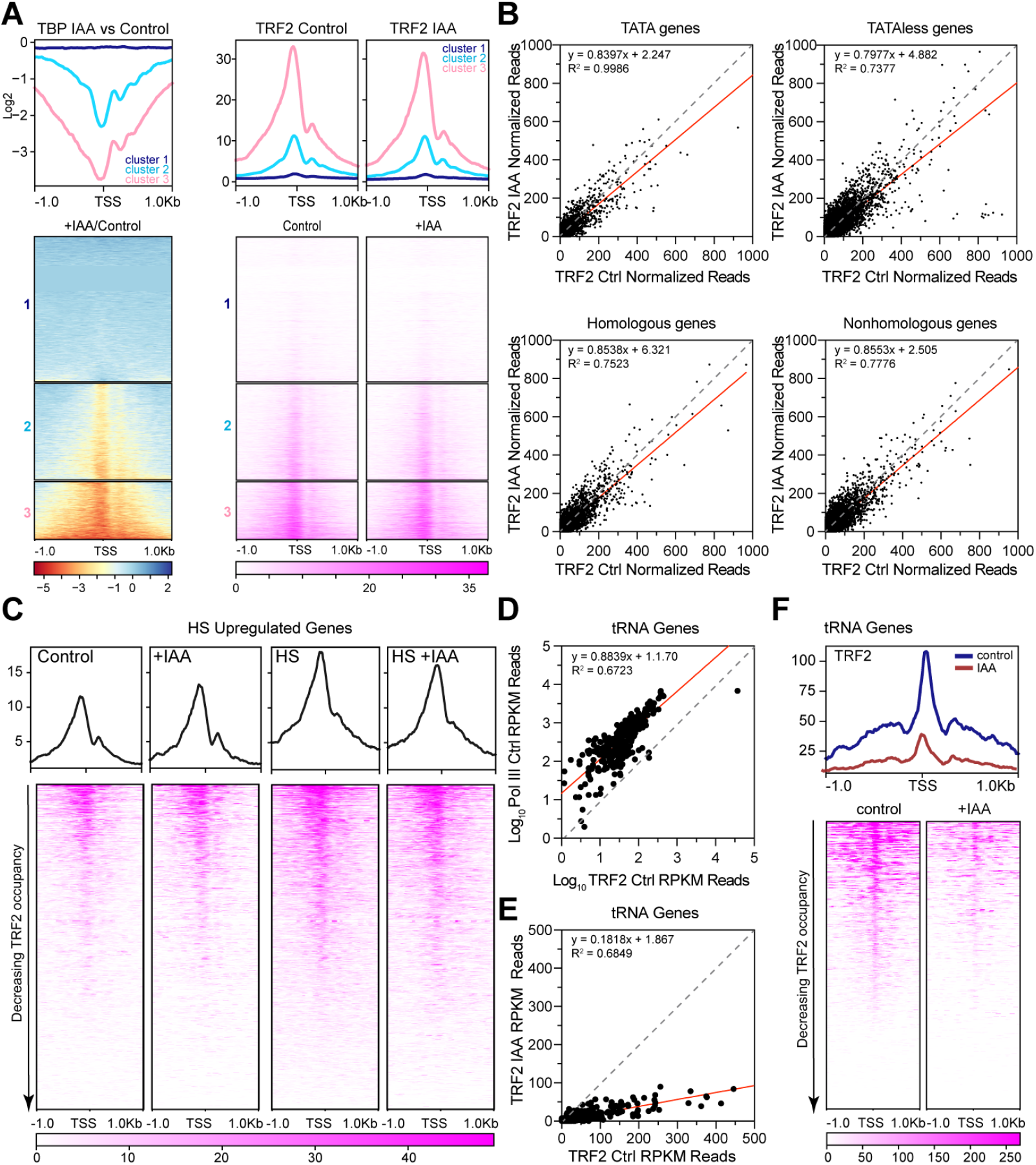
TRF2 binding is TBP-independent in untreated, heat shocked, and retinoic acid treated mESCs. **(A)** k-means clustering with k=3 of the log2IAA/Control of TBP CUT&Tag data (left, same as Figure 1F). TRF2 CUT&Tag average plots and heatmaps by k=3 clustering from left panel in control (middle) and IAA-treated cells (right). **(B)** Scatter plots of log_10_ summed read counts of TRF2 occupancy in control vs IAA-treated C64 mESCs in a 2kb window surrounding the TSS for TATA genes (top left), TATAless genes (top right), yeast homologous genes (bottom left), and non-homologous genes (bottom right). **(C)** Average plots (top) and heatmaps arranged by decreasing TRF2 occupancy (bottom) of TRF2 CUT&Tag in control, IAA-treated (+IAA), heat shocked (HS) and heat shocked +IAA-treated (HS + IAA) C64 mESCs in a 2kb window surrounding the TSS of all genes upregulated by heat shock. **(D)** Scatter plots of log_10_ read counts with RPKM normalization of TRF2 control vs Pol III control C64 mESCs from −50bp to TES of all tRNAs. **(E)** Scatter plots of log_10_ read counts with RPKM normalization of TRF2 control vs TRF2 IAA C64 mESCs from −50bp to TES of all tRNAs. **(F)** Average plots (top) and heatmaps arranged by decreasing TRF2 occupancy (bottom) of TRF2 CUT&Tag in control and IAA-treated (+IAA) C64 mESCs in a 2kb window surrounding the TSS of all tRNA genes with RPKM normalization.

Previous studies have shown that TRF2 activates transcription for a subset of TATA-less but not TATA-containing promoters (82, 84, 85, 87–90). We therefore examined if the presence of the TATA motif affects TRF2 binding in mESCs. We plotted the normalized TRF2 CUT&Tag signal on TATA-containing and TATA-less genes as heatmaps in descending Pol II occupancy levels (Supplemental Figure 8A). Surprisingly, TRF2 binds to both promoter classes at high levels, suggesting that TRF2 does not discriminate between TATA and TATA-less motifs. In contrast to previous studies showing that TRF2 regulates distinct subsets of genes from TBP (91), our data shows that TRF2 binds ubiquitously to all active genes, and that this binding is not mutually exclusive to TBP.

To analyze if TBP depletion affects TRF2 levels on TATA-containing versus TATA-less genes, we determined the normalized read counts for TRF2 with and without IAA treatment at both promoter types and displayed the log values as a scatter plot (Figure 6B Top Panels). Linear regression analysis reveals a strong correlation when comparing control vs. IAA samples for TATA and TATA-less genes (m = 0.8397, R^2^ = 0.9986 and m = 0.7977, R^2^ = 0.7377 respectively). Additionally, we plotted TRF2 for control and IAA conditions as heat maps centered at the TSS for TATA and TATA-less genes (Supplemental Figure 8A). Consistent with the scatter plots, TRF2 remains strongly bound to both promoter types in the presence or absence of TBP.

As a metazoan-specific paralog, TRF2 is not present in yeast (11). We therefore asked if TRF2 binding discriminates between mouse genes homologous to yeast compared to non-homologous genes. Heatmaps of TRF2 CUT&Tag data show that TRF2 binds to both homologous and non-homologous genes (Supplemental Figure 8B), suggesting that mouse genes have evolved to use both TBP and TRF2. For promoters in both homologous and non-homologous genes, we calculated the normalized TRF2 read counts for control vs. IAA samples and displayed the log values as scatter plots (Figure 6B Bottom Panels). Similar to what was seen for TATA-containing and TATA-less genes, we observed little variance between conditions for homologous and non-homologous genes (m = 0.8538, R^2^ = 0.7523 and m = 0.8553, R^2^ = 0.7776 respectively), demonstrating that TRF2 functions independently from TBP regardless of the gene is TATA-containing or TATA-less.

We next examined the genome occupancy of TRF2 during gene induction in control or TBP-depleted cells through the HS response or RA-mediated cell differentiation. Gene browser tracks for HS genes *Hspa1a* and *Hsp90aa1* show low levels of TRF2 binding under control, but increased binding levels after HS (Figure 3B). Surprisingly, we also observe an increase in TRF2 binding throughout the *Hsp90aa1* gene body after HS, though the precise mechanism for this gene body binding remains unclear. We also mapped TRF2 binding after HS in cells depleted of TBP, and observed similar TRF2 levels between HS-only and HS + IAA conditions, with individual replicates showing similar results, suggesting that the increase in TRF2 binding for these genes are independent of TBP (Figure 3B, Supplemental Figure 8C). We next measured TRF2 binding to the top transcriptionally upregulated and downregulated heat shock genes identified from differential gene analysis from Figure 3C. Consistent with the *Hspa1a* and *Hsp90aa1* gene, we detected a slight increase in TRF2 binding at promoters of HS-upregulated genes compared to control samples (Figure 6C, Control vs HS). In contrast, when comparing HS-only and HS + IAA samples, we observe no changes in TRF2 occupancy (Figure 6C), confirming that the increase in TRF2 binding for HS-upregulated genes is indeed independent of TBP. In HS-downregulated genes, we detect similar levels of TRF2 between the control and HS sample, and observe no changes upon TBP depletion (Supplemental Figure 8D).

In RA treatment, gene browser tracks for the *Stra8* gene show relatively low levels of TRF2 under control conditions, consistent with the low levels of TBP and Pol II (Figure 3F). Upon RA treatment, TRF2 binding on the *Stra8* gene increases significantly, tracking with the increase in Pol II and TBP binding (Figure 3F). Under RA + IAA treatment, TRF2 levels remain unchanged compared to RA treatment alone (Figure 3F). As observed with HS gene induction, TRF2 binding to this RA-activated gene (75) appears independent of TBP. We measured TRF2 binding to the top transcriptionally upregulated and downregulated RA genes identified from differential gene analysis from Figure 3G. On average, the RA upregulated genes showed a minor increase in TRF2 binding in RA versus control samples (Supplemental Figure 8F-G). Similarly, this increase in TRF2 occupancy is not perturbed upon TBP depletion when comparing RA and RA + IAA conditions (Supplemental Figure 8F-G). Additionally, downregulated genes show no changes in TRF2 levels when comparing control and RA conditions and when treated with IAA (Supplemental Figure 8E). Taken together with the HS response data, these findings suggest that TRF2 binds to promoters of inducible genes in a TBP-independent manner, which may point to a potential TBP-redundant role for TRF2 in Pol II transcription.

Yeast TBP is required for Pol III transcription, but previous studies have shown that the insect-specific TBP paralog TRF1, and not TBP, is responsible for Pol III transcription in Drosophila (92). Though human TRF2 has been found to be inactive in Pol III transcription *in vitro* (93), we next asked what, if any, role does TRF2 play in Pol III transcription in mESCs. Using our TRF2 CUT&Tag data, we mapped TRF2 binding on tRNA genes (Figure 4A). Surprisingly, we found that TRF2 binds in the same region as TBP in *LeuCAA* and *GlnCTG* genes. We then calculated the TRF2 and Pol III read counts with RPKM normalization for control samples and displayed the log values as a scatter plot for all tRNA genes from −50bp to TES (Figure 6D). We observed a moderate between Pol III levels and TRF2 occupancy (R^2^ = 0.6723). Given that TRF2 binding to Pol II genes is independent of TBP, we next asked how TRF2 binding to Pol III genes is affected when TBP is depleted. We calculated the TRF2 read counts for control and IAA samples and displayed the normalized RPKM values on all tRNA genes (Figure 6E). Strikingly, we observed that most points are well below the diagonal, indicating that TBP depletion leads to decreased TRF2 occupancy on tRNA genes. Additionally, we plotted the TRF2 CUT&Tag RPKM normalized data as a heatmap and average plot in a 2kb window surrounding the TSS of all tRNAs in control and TBP depleted cells (Figure 6F). In agreement with the scatterplot, we see a drastic decrease in TRF2 levels for TBP depleted cells, confirming that TRF2 occupancy at tRNAs is dependent on TBP. Taken together, our analyses show that, though TRF2 binds to promoters of both Pol II- and Pol III-transcribed genes, in the former case, its binding is TBP-independent whereas in the latter, its binding is largely TBP-dependent. Therefore, both TBP and TRF2 function distinctly between these two transcription pathways.

## Discussion

Since the pioneering biochemical and genetic studies on TBP several decades ago, the role of this conserved factor on Pol II transcription has largely remained unquestioned. As an essential gene with roles for at least the three main eukaryotic RNA Polymerases, it has been difficult to define the precise role of TBP in each transcriptional pathway. In this study, we took advantage of a drug inducible degradation system to deplete TBP rapidly and revisit its role in Pol II transcription in mESCs. In contrast to decades of research, we have found that TBP is not required for ongoing Pol II transcription or for induction of silent genes, either via the heat shock response or by cellular differentiation. TBP is, however, required for transcription of tRNA genes by Pol III. The presence of the TBP paralog TRF2 and its binding to promoters of all active genes even in the absence of TBP suggests a potential mechanism for TBP-independent Pol II transcription.

Our surprising results that, in mESCs, TBP is required neither for ongoing transcription nor for activation of silent genes beg the question of the precise role of TBP in Pol II transcription. Previously, we have shown that depletion of TBP in mESCs specifically during mitosis led to impaired reactivation of Pol II genes as cells enter G1 phase (34). One possibility is that, by acting as a mitotic bookmarker, TBP licenses the transcription of all genes right after mitosis such that subsequent transcription events throughout interphase no longer require TBP. How TRF2 functions during mitosis and whether it can act redundantly with TBP during reactivation following mitosis remains unclear. However, given that previous results showing TBP-dependent reactivation following mitosis occur in the presence of TRF2, it is likely that this function may be unique to TBP. Another question that remains to be answered is whether this TBP-independent Pol II transcription is unique to mESCs or if it occurs in other mouse cell types. Since TRF2 expression is fairly widespread across mouse cell types (94), we predict that this phenomenon is more general, though whether other mammalian cells, such as human cells, require TBP for Pol II transcription remains to be seen. Results from this study provide new context for the role of TBP in Pol II transcription, specifically in mammalian systems.

Historically, TBP has been shown to be important for transcription initiation from TATA-containing promoters. Defects in the TBP DNA binding surface in yeast have been shown to be detrimental for transcription initiation at TATA-containing promoters, but not for TATA-less containing promoters *in vivo* (40). However, depletion of TBP from the yeast nucleus has been shown to completely downregulate global transcription in both TATA and TATA-less genes (18). It is apparent that TBP is still required for transcription initiation and PIC formation in budding yeast in both TATA-less and TATA-containing genes, even though the role of TBP at TATA-less promoters and how the PIC forms at TATA-less promoters remain to be elucidated. *In vitro* studies have shown that yeast and human TBP can support basal transcription without the other TFIID members from TATA-containing promoters, but not from TATA-less promoters (44, 95, 96). Mutating the TATA box of a TATA-containing DNA template abolishes binding of yeast TBP, which leads to a subsequent decrease of transcription *in vitro* (14). However, TATA-less genes still require TBP for transcription *in vitro*, since immunodepletion of human TBP from HeLa cell extracts decreased transcription from both TATA-containing and TATA-less promoters *in vitro* (44). Surprisingly, we observed that Pol II transcription of both TATA-containing and TATA-less genes was largely unaffected by the depletion of TBP in mESCs. Even for genes homologous to budding yeast, TBP depletion did not lead to decreased transcription of these genes in mESCs, suggesting that homologous genes are not intrinsically more TBP-dependent than non-homologous genes. These findings suggest that the mechanisms of transcription initiation in mammals have evolved to not only be TBP-independent but also independent of the presence of the TATA-box on gene promoters.

From our data, we observed that genes enriched with paused Pol II were activated by heat shock or RA treatment even in the absence of TBP. Many studies have focused on the role of TBP in transcription initiation and PIC formation, but its roles in downstream processes like transcription pausing, elongation, and pause release are not as thoroughly explored. Early *in vitro* experiments found that after Pol II escapes the promoter, TFIID, TFIIA, TFIIH, and TFIIE remain on the promoter as a scaffold that could allow for subsequent Pol II loading (97). For heat shock genes in yeast, PIC components are enriched at the gene promoter, even if the gene is not being actively transcribed. For example, TBP, TFIIB, TFIID, and TFIIH occupy heat shock genes in non-heat shock conditions at different levels, despite the lack of transcription of these heat shock genes in yeast (18, 98). Similarly, from our data we observe TBP on heat shock genes in control conditions when these genes are not being actively transcribed (Figure 3B). When paused genes are activated, GTF occupancy levels tend to increase on the promoters of the activated genes (98). Our data show a similar increase in TBP binding on the promoters of heat shock and RA-induced genes upon heat shock and RA treatment respectively (Figure 3B, 3F), which suggest a role for TBP in the transcription activation of paused genes. However, we observe the upregulation of heat shock and RA-induced genes even when TBP is depleted, which suggests that TBP is not required for the recruitment and loading of Pol II for subsequent rounds of transcription after Pol II pause release. One potential explanation to this observation is that paused Pol II intrinsically prevents nucleosome formation over promoters, which alone could be sufficient for subsequent Pol II loading (99). Another possibility is that a PIC scaffold could form without TBP at paused gene promoters, thus eliminating the need for TBP in the transcription activation of paused genes (97), though the presence of such a scaffold is still in debate. Whether a PIC without TBP is formed on promoters of paused genes or not, we observe that transcription activation of paused genes by heat shock and retinoic acid differentiation occurs in a TBP-independent manner.

Though the core members of the PIC are highly conserved, gene duplication events and subsequent specialization throughout evolution have led to the emergence of PIC paralogs that contribute to non-canonical initiation complexes. For example, TBP associated factors (TAFs) that make up TFIID have acquired paralogs that are often expressed in a cell-type specific manner (100–103). In fission yeast, the TFIID complex could be composed of either spTAF5 or spTAF5b, two TAF5 homologues that are expressed (104). Multiple TAF paralogues have also been reported in *C. elegans, D. melanogaster* and humans (105–107). Tissue specific TAF paralogs such as TAF3 has been implicated in embryonic stem cell lineage commitment, TAF4b in maintenance of B lymphocytes located in ovaries and testes, and TAF9B, an orphan TAF that does not associate to the canonical TFIID complex, in motor neuron differentiation, further diversifying the already complex transcriptional repertoire (108–111). Historically, TBP has been thought to be a universal eukaryotic transcription factor due to its high conservation across organisms, its presence in all three eukaryotic polymerases and the embryonic lethality that accompanies homozygous TBP knock-out (23, 112). However, the discovery of TBP paralogs with their own unique PIC members further partitioned the genome, challenging the universal role of TBP. In fact, the TBP paralog TRF3 has been proposed to functionally replace TBP in *Xenopus* oocyte transcription (26). Here, we show that depletion of TBP in mESC does not directly affect RNA Pol II transcription, and propose that a non-canonical PIC containing TRF2 may facilitate Pol II transcription in its place (Figure 1D, Figure 5, Figure 6).

TRF2 (also known as TLF) is a more distant TBP paralog compared to the other paralogs, and is found in all multicellular organisms except *S. cerevisiae*. TRF2 shares only ~40% amino acid identity with the core domain of TBP (113), and in *Drosophila*, TRF2 fails to bind to DNA that contains the canonical TATA box. Instead, TRF2 from various species such as *Drosophila*, worms, frogs, fish, and flies has been proposed to activate transcription for TATA-less promoters *in vitro* (11, 30, 83, 89, 113). For example, a previous genome-wide occupancy profile of *Drosophila* TRF2 has shown that about 80% of TRF2 binding sites do not overlap with TBP (88). Surprisingly, our CUT&Tag results show that TRF2 binds strongly to both TATA and TATA-less promoter types in mESCs (Supplemental Figure 8A). We report that TRF2 binds to most active genes, similarly to TBP, and that the levels of TRF2 largely correlate with levels of TBP and Pol II (Figure 5D-E). One possible explanation for various TRF2 functions is the variability in the TRF2 core domain across species. Unlike TBP, whose core domain is over 80% conserved across species, TRF2’s core domain shares only 40-45% amino acid conservation between metazoans (11, 114). We also show that TRF2 physically interacts with Pol II (Figure 5B) and experiences a similar increase in promoter occupancy as TBP during gene induction via HS and RA (Figure 3B, Figure 3F). The binding of TRF2 also seems to be independent of TBP as depletion of TBP does not affect TRF2 binding nor its increase in occupancy during gene activation (Figure 6C-D). Although TRF2 has been historically seen to govern specific subsets of genes distinct from TBP in certain species, our results show that TRF2 in mESCs have the potential to substitute for TBP during transcription initiation of all mouse Pol II-transcribed genes.

The TBP-independent Pol II transcription in mESCs contrasts strikingly with the acute impairment of Pol III transcription in the absence of TBP. Early *in vitro* studies using purified factors from cultured human KB cells showed that TBP is a general component of Pol III transcription (22). In addition, TBP inactivation in budding yeast with temperature-sensitive alleles of *TBP* led to a rapid decrease of Pol III transcription *in vivo* (2). Our results showing that TBP is required for Pol III activity in mESCs therefore serve two purposes: first, it provides confirmation that our acute TBP depletion system in mESCs has the expected consequences, at least when it comes to Pol III; and second, it provides a sharp emphasis on the lack of effect on Pol II transcription when TBP is depleted. Though we confirm that mouse TBP is required for Pol III-mediated transcription, this result is also surprising in a way given that previous studies in Drosophila has shown that TRF1, a widely expressed TBP paralog, was part of the Pol III pre-initiation complex instead of TBP (92). Importantly, thus far, Drosophila TRF1 has been the only TBP paralog shown to be important for Pol III transcription. Therefore, our result showing that mouse TRF2 binds to tRNA genes is the first evidence of a mammalian TBP paralog playing a role in Pol III transcription. Indeed, human TRF2 was not able to functionally replace TBP in transcription by Pol III *in vitro* (93), and the oocyte-specific mouse TRF3 does not interact with Pol III PIC subunits, suggesting that even in oocytes, mouse TRF3 does not play a role in Pol III-mediated transcription (33). Despite our observation that mouse TRF2 binds to tRNA genes, its precise role in Pol III transcription remains unclear. In fact, our observation that TRF2 levels decrease upon TBP depletion across tRNAs (Figure 6E-F) suggests that its function is downstream of TBP in Pol III transcription, which is in contrast to the TBP-independent binding of TRF2 across Pol II-transcribed genes. This distinct specificity lends further support for TRF2 enabling TBP-independent Pol II-transcription in mESCs.

## Materials and Methods

### Cell culture

For all experiments, the mouse ES cell line C64 was used, which is a CRISPR genetically modified JM8.N4 (RRID: CVCL_J962) cell line obtained as previously described (Teves et al. 2018). ES cells were cultured on 0.1% gelatin-coated plates in ESC media Knockout D-MEM (Invitrogen, Waltham, MA) with 15% FBS, 0.1 mM MEMnon-essential amino acids, 2 mM GlutaMAX, 0.1 mM 2-mercaptoethanol (Sigma) and 1000 units/ml of ESGRO (Chem-icon). ES cells are fed daily, cultured at 37 °C in a 5% CO2 incubator, and passaged every two days by trypsinization. For endogenously-tagged mAID-TBP cells, TBP degradation was performed by addition of indole-3-acetic acid (IAA) at 500 μM final concentration to a confluent plate of cells for 6 hours. Heat Shock was performed at 42 °C in a 5% CO2 incubator for 30 minutes and Heat Shock Recovery cells were shifted back to 37 °C in a 5% CO2 incubator for 30 minutes after Heat Shock. For Heat Shock and IAA treatment, cells were first treated with 6 hours of auxin followed by an additional 30 minutes to 1 hour due to heat shock and recovery before being collected. Retinoic Acid (Sigma-Aldrich R2625-100MG) treatment was performed at 37 °C in a 5% CO2 incubator at 0.25 μM for 16 hours. Cells treated for Retinoic Acid and IAA were incubated for 16 hours at 0.25 μM and 500 μM, respectively.

### NET-seq

Cell fractionation was performed as described in Mayer and Churchman 2016 (78). Mouse ES cells were scraped, collected by centrifugation, and washed twice with ice-cold PBS buffer. Approximately 10 to 15 million cells were used in each fractionation. Drosophila S2 cells were added to each fractionation (3% by cell count) as spike-in for downstream analyses. Cells were lysed with cytoplasmic lysis buffer (0.15% v/v NP-40, 10 mM Tris-HCl pH 7.0, 150 mM NaCl, 25 uM α-amanitin, 10 U SUPERase. In, 1X protease inhibitor mix) by pipetting the sample up and down ten times, followed by incubation on ice for 7 minutes. Cell lysates were layered onto 500 μL of sucrose buffer (10 mM Tris-HCl pH 7.0, 150 mM NaCl, 25% w/v sucrose, 25 uM α-amanitin, 20 U SUPERase. In, 1X protease inhibitor mix). Nuclei was collected by centrifugation, and the supernatant (cytoplasmic fraction) was collected for the Western blot control sample. The nuclei were washed twice with nuclei wash buffer (0.1% v/v Triton X-100 1 mM EDTA, 1X PBS, 25 uM α-amanitin, 40 U SUPERase. In, 1X protease inhibitor mix). Nuclei were resuspended in glycerol buffer (20 mM Tris-HCl pH 8.0, 75 mM NaCl, 0.5 mM EDTA, 50% v/v glycerol, 0.85 mM DTT, 25 uM a-amanitin, 10 U SUPERase, 1X protease inhibitor mix) and lysed in nuclei lysis buffer (1% v/v NP-40, 20 mM HEPES pH 7.5, 300 mM NaCl, 1 M urea, 0.2 mM EDTA, 1 mM DTT, 25 uM a-amanitin, 10 U SUPERase, and 1X protease inhibitor mix) by pulsed vortexing and incubating on ice for 2 minutes. Chromatin was collected by centrifugation, and the supernatant (nucleoplasmic fraction) was collected for the Western blot control sample. RNA from the insoluble chromatin was extracted with Trizol for library preparation. For the Western blot analysis of the control sample, chromatin was solubilized with 1U MNase and incubated at 37 °C for 10 minutes for downstream Western blot analyses.

Barcoded DNA linkers were riboadenylated in-house as followed in Song et al (115). Briefly, adenylation reactions were performed in a reaction mixture (10 μL) containing 0.8 ug of barcoded DNA linker, 1X T4 RNA ligase buffer, 2 mM ATP, 35% PEG, and 300 U of T4 RNA Ligase 1, incubated at 37 °C for 8 hours, followed by 15 minutes of heat inactivation at 65 °C. Riboadenylated barcoded DNA linkers were stored at −80 °C until use.

NET-seq library preparation was performed as described in Mayer and Churchman 2016 (78). RNA samples were denatured for 2 minutes at 80 C before DNA linker ligation. Approximately 1 ug of denatured RNA was ligated with barcoded DNA linker in a reaction mixture containing 1X T4 RNA ligase buffer, 10% DMSO, 20% PEG, 200 U of Truncated T4 RNA ligase 2, and 0.91 ug of the in-house riboadenylated barcode DNA linker, incubated at 37 °C for 3 hours. Ligations were quenched with the addition of 0.7 μL of 0.5 M EDTA. To fragment the RNA, 2X alkaline fragmentation solution (0.012 M Na_2_CO_3_, 0.088 M NaHCO_3_) was added to the reaction mix and incubated at 95 °C for 75 minutes. RNA precipitation solution (0.32 M sodium acetate, pH 5.5), isopropanol, and co-precipitant GlycoBlue were added to each sample and left at −80 °C overnight to purify the fragmented RNA. The fragmented RNA samples were pelleted by centrifugation, resuspended with RNAse-free H_2_O, and briefly denatured with 2X TBU denaturing sample buffer. The samples were loaded onto a Novex denaturing 15% poly-acrylamide TBE-urea gel (Invitrogen) and ran according to the manufacturer’s instructions. The gel was stained with SYBR Gold (Invitrogen) and the region between 35 and 100 nt was excised. The size-selected RNA was extracted from the gel and purified by isopropanol as described in Mayer and Churchman 2016 (78).

Reverse transcription of the fragmented RNA was performed using the SuperScript™ III First-Strand Synthesis System Kit (Thermofisher) as described in Mayer and Churchman 2016 (78). Briefly, gel-purified RNA fragments were resuspended in RNAse-free dH_2_O and mixed with RT reaction mix at a final concentration of 250 mM Tris, 375 mM KCl, 15 mM MgCl_2_, 0.5 mM dNTPs, and 0.3 uM reverse primer. Each sample mixture (14.6 μL) was incubated for 2 minutes at 80 °C and chilled on ice for 3 minutes. RNA was then reverse transcribed with 1.3 μL of SUPERase, DTT mix (10 U SUPERase. In, 0.06 M DTT) 160 U of SuperScript III RT, and incubated at 48 °C for 30 minutes. The RT reaction was quenched with 1.8 μL of 1 M NaOH, incubated at 98 °C for 20 minutes, and neutralized with 1.8 μL of 1 M HCl and chilled on ice. The cDNA samples were briefly denatured with 2X TBU denaturing sample buffer, loaded onto a Novex denaturing 10% poly-acrylamide TBE-urea gel (Invitrogen) and ran according to the manufacturer’s instructions. The gel was stained with SYBR Gold (Invitrogen) and the region between 85 and 160 nt was excised. The size-selected cDNA was extracted from the gel and purified by isopropanol as described in Mayer and Churchman 2016 (78). The first strand cDNA was then resuspended in RNAse-free H2O and circularized using the CircLigase ssDNA ligase kit (Mandel Scientific) according to the manufacturer’s instructions.

cDNA containing sequences of a subset of sequenced snRNAs, snoRNAs, rRNAs and tRNAs were specifically depleted using biotinylated DNA oligos as described in Mayer and Churchman 2016 (78). Briefly, biotinylated DNA oligos were added to circular single-stranded DNA and subjected to subtractive hybridization in a thermal cycler. Streptavidin-coupled magnetic beads were then added to the depletion reaction to pull down the heavily sequenced mature RNAs. The circular, depleted cDNA was purified with isopropanol, 0.375 M NaCl, and GlycoBlue as a co-precipitant overnight at −20 °C.

Circular cDNA was amplified by PCR using Phusion High-Fidelity enzyme (NEB) according to the manufacturer’s instructions. PCR was carried out with an initial 30s denaturation at 98 °C, followed by 6 cycles of 10s denaturation at 98 °C, 10s annealing at 60 °C, and 5s extension at 72 °C. The PCR products were loaded onto a Novex denaturing 8% poly-acrylamide TBE-urea gel (Invitrogen) and ran according to the manufacturer’s instructions. The gel was stained with SYBR Gold (Invitrogen) and the ~150 nt band was excised. The size-selected NET-seq libraries were extracted from the gel with DNA soaking buffer (0.01 M Tris-HCl pH 8.0, 0.3 M NaCl, 1 mM EDTA) shaking at 1500 rpm overnight in a Thermomixer at room temperature. The NET-seq libraries were purified by isopropanol precipitation overnight at −20 °C and resuspended with 10 mM Tris-HCl pH 8.0. Sequencing was performed at the UBC Biomedical Research Centre with 55 bp single-end reads.

### Processing and alignment of NET-seq reads

NET-seq data was processed as described in Mayer et al. 2015 (38). Reads were trimmed and aligned using STAR (v. 2.4.0). Reverse transcription mispriming events, PCR duplication events, and splicing intermediates were removed using custom Python scripts provided by the Churchman group (https://github.com/churchmanlab). The bam files of biological replicates were merged and normalized by subsampling each sample by the number of aligned Drosophila reads across all samples using Samtools. Coverage files for gene plots were generated from normalized bam files with a custom Python script provided by the Churchman group (https://github.com/churchmanlab) and visualized by IGV. Read counts, heatmaps, and TSS plots were performed using DeepTools and BedTools. Read counts were plotted and analyzed by linear regression using GraphPad Prism.

### CUT&Tag Protocol

Concanavalin A-coated beads (Cedarlane Labs BP531-3ML) were prepared as previously described (37), using 10 μL of beads per sample. Cells were harvested at room temperature and 100,000 mESCs were used per sample. Cells were pelleted at room temperature and then washed using 1 volume of Wash Buffer twice (20 mM HEPES pH 7.5, 150 mM NaCl, 0.5 mM Spermidine in dH2O with 1X Roche Complete Protease Inhibitor EDTA-Free tablet). Cells attached to beads were then resuspended in Antibody Buffer (2 mM EDTA, 0.101% BSA in Dig-wash buffer and chill on ice before use) at a final volume of 50 μL per sample with 1μg of primary antibody was added to each sample and incubated 4 °C overnight on a nutator. Antibodies used include TBP (Abcam ab51841), RNA Pol II (Cell Signalling Technologies D1G3K), TRF2 (In house antibody gift from Dr. László Tora), IgG (Abcam ab46540), K27 (Cell Signalling Technologies C36B11). Cells were collected using a magnetic rack, and 100 μL secondary antibody mixed in a 1:100 ratio in Dig-wash Buffer (0.05% digitonin in Wash buffer) was added to each sample mixed at room temperature for 45 minutes. Cells were collected as before and washed three times with 0.8-1 mL Dig-wash Buffer. The Henikoff lab pA-Tn5 adapter complex or the commercial Epicyper pA-Tn5 (Epicypher EP151117) was added to a final concentration of 1:250 or 1:20, respectively, per sample in Dig-300 Buffer (20 mM HEPES pH 7.5, 300 mM NaCl, 0.5 mM Spermidine, 0.01% Digitonin in dH2O). Samples were washed and the tagmentation reaction was performed in Tagmentation Buffer (10 mM MgCl_2_ in Dig-300 Buffer) for 1 hour in a 37 °C water bath. Tagmentation was stopped and DNA fragments were solubilized by adding 10 μL 0.5 M EDTA, 3 μL 10% SDS and 2.5 μL 20 mg/mL Proteinase K to each sample. Samples were then vortexed at maximum speed for 5s and incubated in a 37 °C water bath overnight to digest. DNA was extracted using phenol chloroform, and library preparation was performed as previously described.

### CUT&Tag Analysis

Reads were mapped on mm10 genome build using Bowtie2 with the following parameters: --no-unal --local --very-sensitive-local --no-discordant --no-mixed --contain --overlap --dovetail --phred33 –I 10 –X 9999. PCR duplicate reads were kept as these sites may represent real sites from adapter insertion from Tn5 as per recommendation from the Henikoff lab. A normalization factor was determined from *E. coli* (normalized to the control) alignment from Bowtie2 mapping and used to scale samples during generation of bigwig files. Samples without spike-in control include Pol III CUT&Tag. For these samples, normalization was done by RPKM. Downstream analyses, heatmaps, TSS plots, Gene plots, k-means clustering were performed using IGV, DeepTools and BedTools suite. ComputeMatrix from deeptools was done using binsize 10.

### Differential Gene Analysis using edgeR bioconductor

Scatter plots of differential gene analysis were performed using the DGE (Differential Gene Expression) tool of edgeR bioconductor package. Starting bam files were used for all samples. Read counts were obtained using featureCounts command with paired end specificity for the entire gene and pseudogenes were removed. Reads were then imported into R Studio and analyzed using DGE tool following the bioconductor manual using default parameters (ie. genes are 5% FDR). First, DGEList was used to specify reads and gene names, followed by gene annotations (optional) then filtering and normalization by TMM (default option). Read counts below 20 were filtered using filterByExpr(min.count = 20). Once reads are normalized, the design matrix was built and dispersion was estimated to determine biological variation. The command glmFit was used to determine differentially expressed peaks/genes (DEG) and the command topTags was used to show the top genes. Scatter plots were generated using the plotMD function and significant genes were obtained along with the raw values for downstream analysis such as heatmap and table generation.

### Gene Ontology

Gene ontology was done on gene sets identified to be significant from differential gene analysis. The top HS and RA upregulated genes were inputted into http://geneontology.org/, filtered by molecular function for the Mus Musculus genome to obtain GO terms.

### Scatter Plot Read Counts Analysis

Read counts for scatter plots were obtained from RNA Pol II bam files using the bedtools multicov command. Read counts were then normalized to the *E. coli* scaling factor (*E. coli* reads in treatment versus control) or to RPKM values and plotted as is or Log_10_ transformed before being plotted. Linear regression was then performed to obtain the slope and R^2^ value.

### Pausing Index Analysis and Enhancer Analysis

Pausing Index and Enhancer Analysis were both done through the NRSA (Nascent RNA Sequencing Analysis) suite. RNA Pol II bam files were obtained from CUT&Tag data and the NRSA suite pause_PROseq.pl command was used to obtain the pausing index box plot using the default values with -in1 as condition 1 and -in2 as condition 2. Box represents the interquartile range (middle 50% of the data), and the whiskers represent the upper and lower range (25% of the remainder). Average pausing index was obtained from averaging the raw pausing index of each gene obtained from pause_PROseq.pl, taking the median value of all averaged genes, and plotting in R studio using boxplot function. Pausing Index for HS and RA genes were done using the top upregulated genes extracted from edgeR analysis. Enhancer analysis was obtained using the eRNA.pl command for RNA Pol II bam files with default values -pri 1 -dir 0 -lk pp -cf 0.001 to obtain the RNA Pol II signal around the center of enhancers for all samples.

### Immunofluorescence

Cells were grown in coverslips (Azn Scientific #ES0117650) pre-washed in 70% ethanol and coated in gelatin in tissue culture-treated 6-well plates. After indicated treatments, cells washed with 1 mL of PBS and fixed 4% Paraformaldehyde (UBC Chemical Stores #OR683105) for 15 minutes. After fixation, cells were washed and permeabilized with 1 mL 0.025% Triton-X for 5 minutes, followed by 2 10-minute washes with PBS. Samples were blocked by nutating in PBS 5% BSA for 30 minutes. Primary antibodies were diluted at 1:100 in PBS, and added to cells in coverslips for 1 hour. Samples are washed twice with PBS with 5 minutes for each wash. Coverslips were then incubated in secondary antibody 1:100 for 1 hour, and washed with PBS twice. Samples are then incubated with DAPI (300 nM in PBS) for 5 minutes and washed twice with PBS with 5 minutes for each wash. Coverslips were assembled using Vectashield mounting medium (BioLynx #VECTH1000). Fluorescent images were collected using the Leica DMI6000B inverted fluorescence microscope. Quantification was done through Fiji (116).

### CoIP

Cells were grown to 80% confluency on a 10 cm gelatinized plate. Cells are then washed 2X with PBS and scraped using ice-cold PBS 0.2 mM PMSF (Sigma-Aldrich #11359061001) 0.001% Aprotonin (Sigma-Aldrich #A6279-10ML) solution. Cells are pelleted and resuspended in 1 mL of Lysis Buffer (200 mM NaCl, 25 mM HEPES, 1 mM MgCl_2_, 0.2 mM EDTA, 0.5% NP-40, 1X Roche complete inhibitor, 0.2 mM PMSF, 0.001% Aprotonin, 1 mM DTT, 1 mM benzamidine). Solution is then passed through a 25G needle 10X and rocked at 4 °C for 30 minutes. Samples were spun at max speed at 4 °C for 30 minutes. Supernatant is transferred to a new tube and pellet was washed several times with dH2O. Pellet was then digested as much as possible using 2 U of MNase in 1X MNase digestion buffer by nutating at 37 °C for 10 minutes followed by shearing with a 25G needle. Digested pellet is centrifuged at max speed and the new supernatant is added to the previous supernatant. Residual pellet is resuspended in 100 μL of 1X SDS. Lysates were then pre-cleared using 20 μL/sample of Protein A+G Magna ChIP beads and incubated on an end-over-end rotator for 2 hours at 4 °C. Samples are then placed on a magnetic rack and aliquoted (50 μL for input and divide the rest depending on the number of IPs). Then, 50 μL of 2X SDS was added to the input for a 100 μL 1X SDS final solution. Pull down was done by adding 1-3 μg of primary antibody to the samples and incubated on an end-over-end rotator for 2 hours at 4 °C. Another set of 40 μL beads is pre-cleared using Lysis Buffer for 2 hours at 4 °C. Pre-cleared beads were pelleted and resuspended in 50 μL/sample of Lysis Buffer to add to lysate. Samples are incubated on an end-over-end rotator overnight at 4 °C. The following day samples are placed on a magnetic rack to withdraw the liquid. Beads are then washed 4X using 600 μL Lysis Buffer. Beads are resuspended in 100 μL of 1X SDS and boiled at 100 C for 10 minutes followed by max speed centrifugation for 5 minutes. For Western blot analysis, 20 μL of sample is loaded on to a 8% SDS-PAGE for Pol II and 4% for TBP/TRF2. Antibody concentrations used for Western Blots were 1:3000 primary and 1:10000 secondary. Antibodies used include: TBP (Abcam ab51841), Pol II (In house antibody from Dr. Robert Tjian), TRF2 (In house antibody gift from Dr. László Tora)

## Author Contributions

J.Z.J.K. and T.F.N contributed equally to this work. J.Z.J.K., T.F.N, and S.S.T. conceived and designed the study. J.Z.J.K., T.F.N, and M.A.B. developed the methodology. J.Z.J.K., T.F.N, M.A.B, R.M.P, and J.C. acquired the data, and performed analyses. J.Z.J.K., T.F.N, and S.S.T wrote and revised the manuscript, with critical comments and editing from M.A.B, R.M.P and J.C. The study was supervised by S.S.T.

## Acknowledgements

We thank Dr. Steven Henikoff for providing the pA-Tn5 enzyme, and D. Laszlo Tora for providing the anti-TRF2 antibody. We thank R. Vander Werff and T. Stach (BRC-seq, UBC) for Illumina sequencing and S. Flibotte (LSI Bioinformatics facility, UBC) for implementation of NET-seq analyses. For insightful comments on the manuscript, we thank Drs. Annie Ciernia and Ethan Greenblatt. M.A.B is a postdoctoral fellow supported by the Sigrid Jusélius Foundation. S.S.T is a Wall Scholar at the Peter Wall Institute, and is supported by the Michael Smith Foundation for Health Research. This work was supported by the Canadian Institutes for Health Research Project Grant award to S.S.T. (PJT-162289) and by the National Sciences and Engineering Research Council Discovery Grant award to S.S.T. (RGPIN-2020-06106).

