## Supplemental Figures for "A TBP-independent mechanism for RNA Polymerase II transcription"

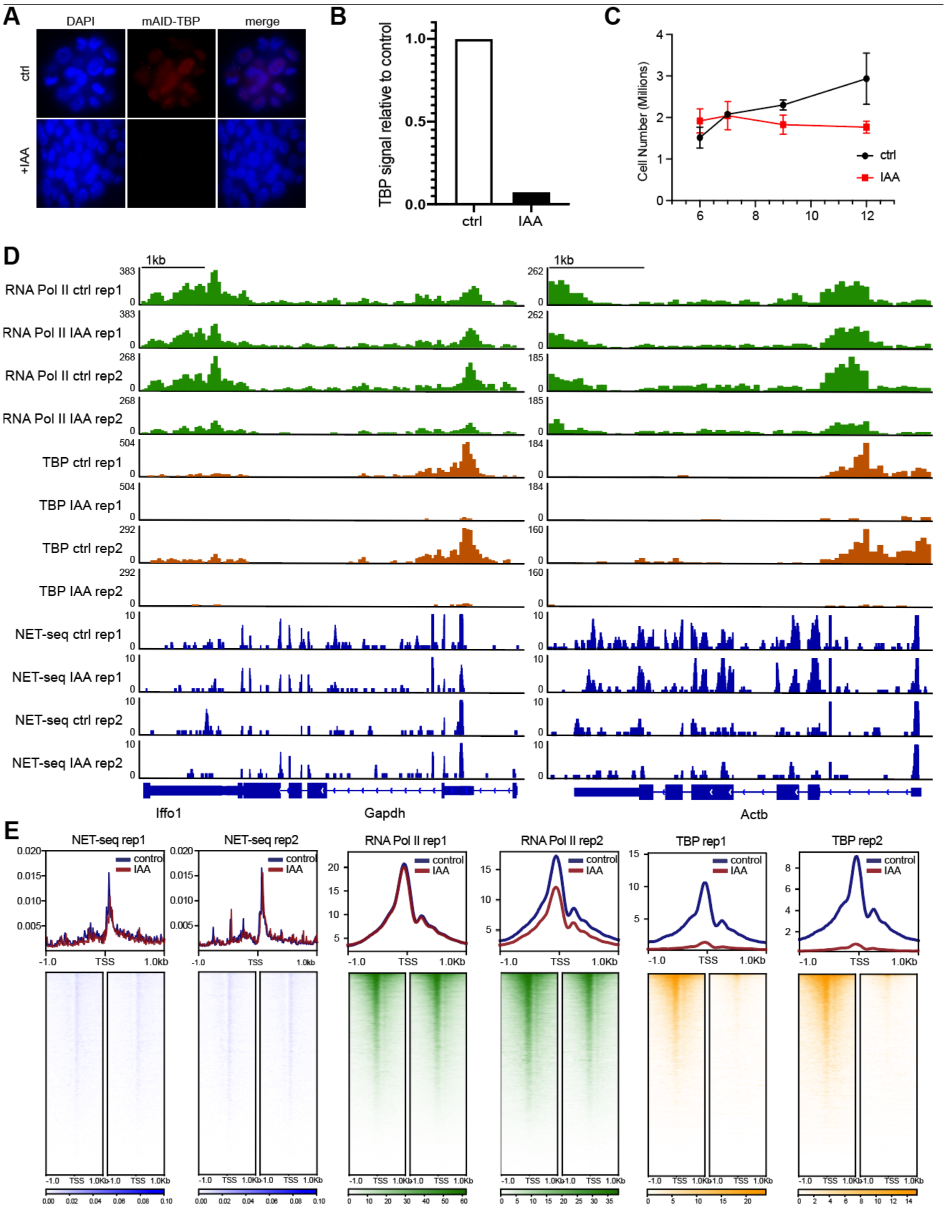


**Supplemental Figure 1. Replicate analysis of TBP and Pol II CUT&Tag, and NET-seq data.**

**(A)** Immunofluorescence using α-TBP antibody of C64 mESCs with endogenous mAID-TBP under control conditions (top) and after 6 hours of IAA treatment (bottom). **(B)** Quantification of fluorescence signal from (A). **(C)** Cell growth curve of C64 mESCs from 6 hours to 12 hours of treatment with DMSO (ctrl, black) or IAA (IAA, red). **(D)** Gene browser tracks at *Gapdh* (left) and *Actb* (right) for CUT&Tag analyses of TBP (orange), TRF2 (magenta) and Pol II (green), and strand-specific reads from NET-seq data (blue) of two biological replicates of control (Con) or IAA-treated (IAA) C64 mESCs. **(E)** Genome-wide average plots (top) and heatmaps arranged by decreasing Pol II occupancy (bottom) of NET-seq (blue), Pol II CUT&Tag (green), and TBP CUT&Tag (yellow) in a 2kb window surrounding the TSS of all genes of two biological replicates of control and IAA treated C64 mESCs.

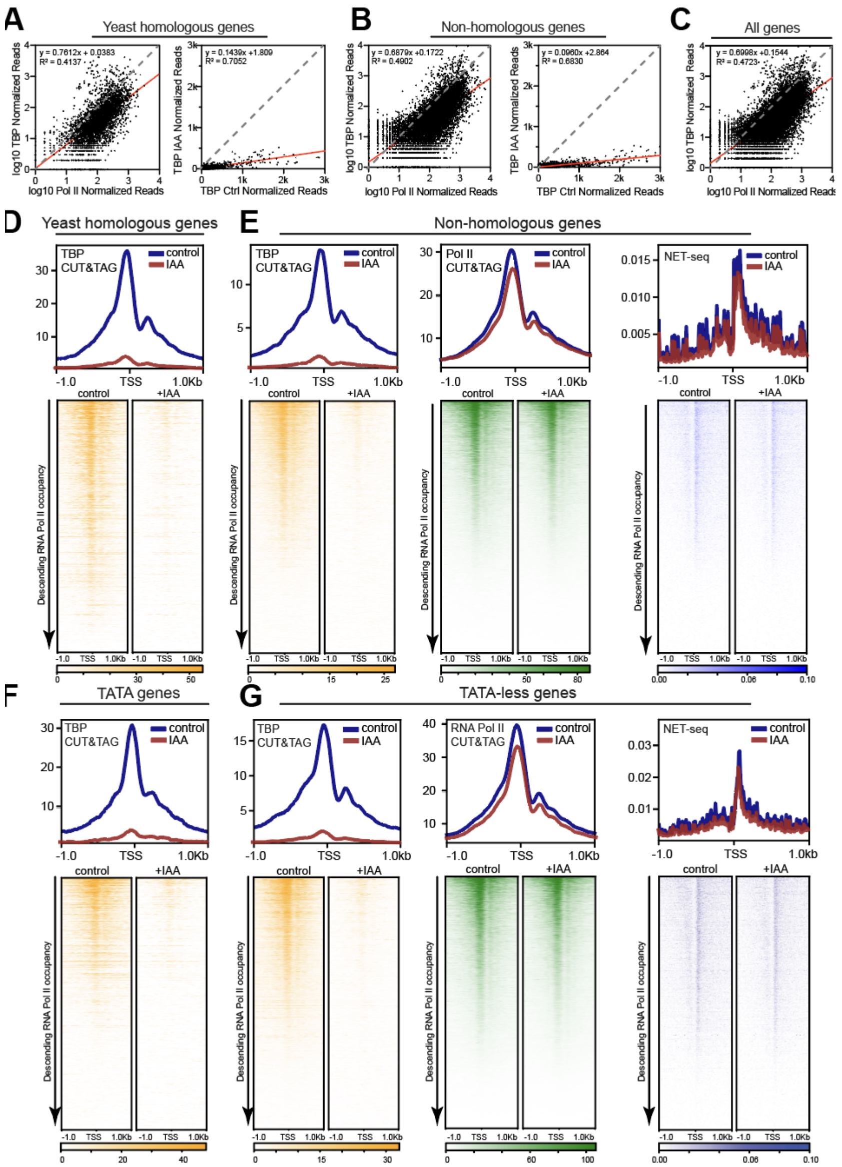


**Supplemental Figure 2. TBP is not required for transcription of specific subsets of genes.**

**(A-B)** Scatter plots of $log_{10}$ summed read counts of Pol II vs. TBP occupancy in control C64 mESCs (left) or TBP occupancy in control vs IAA-treated C64 mESCs (right) in a 1kb window surrounding the TSS for yeast homologous genes (A) and non-homologous genes (B). **(C)** Scatter plots of $log_{10}$ summed read counts of Pol II vs. TBP occupancy in control C64 mESCs in a 1kb window surrounding the TSS for all genes. **(D,F)** Average plots (top) and heatmaps arranged by decreasing Pol II occupancy (bottom) of TBP CUT&Tag (yellow) in a 2kb window surrounding the TSS of yeast homologous (D) and TATA-containing genes (F). **(E,G)** Average plots (top) and heatmaps arranged by decreasing Pol II occupancy (bottom) of TBP CUT&Tag (yellow), Pol II CUT&Tag (green), and NET-seq (blue) in a 2kb window surrounding the TSS of non-homologous **(E)** and TATA-less genes **(G)**.

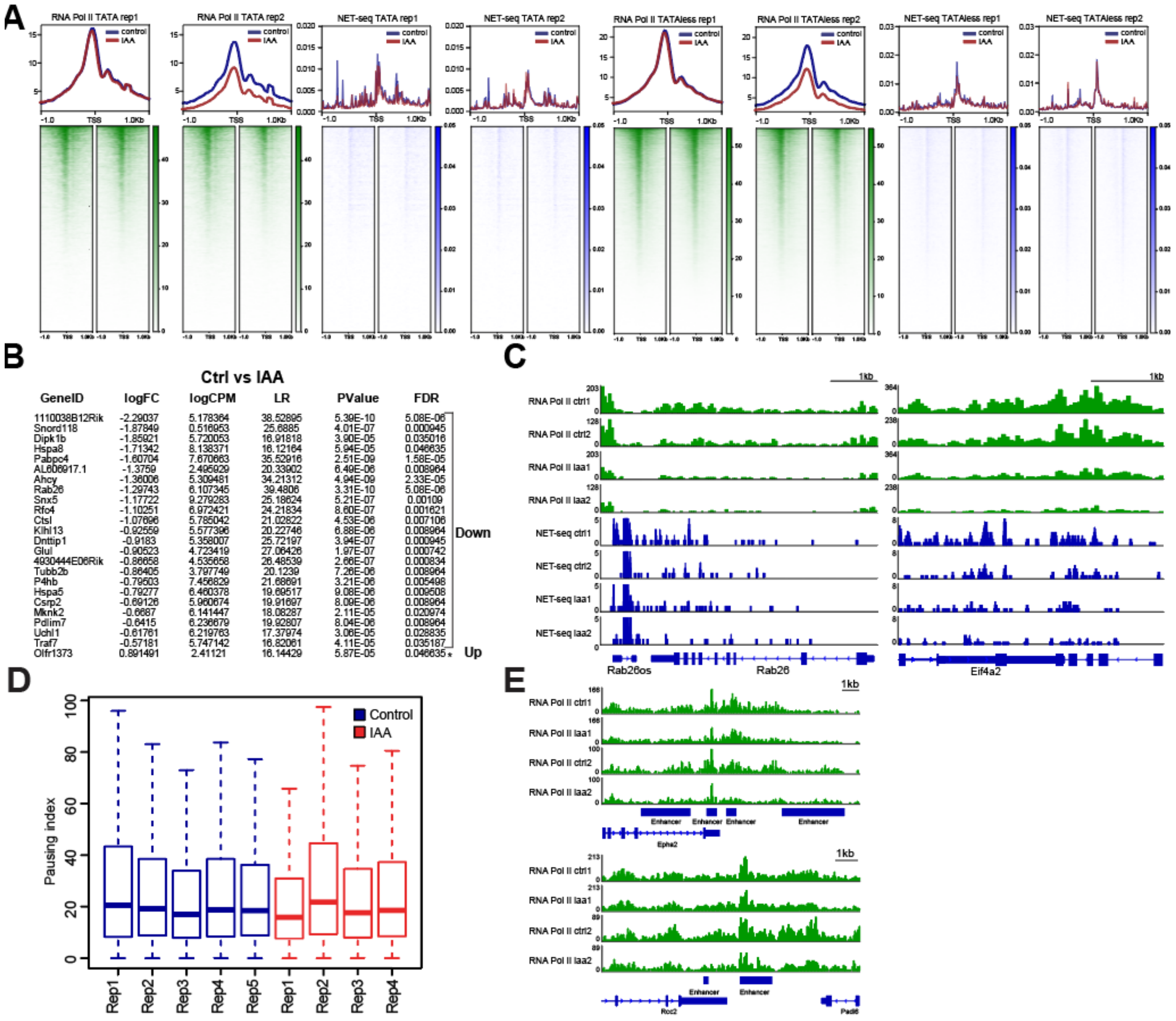


| GeneID | logFC | logCPM | LR | PValue | FDR |
|---|---|---|---|---|---|
| 1110038B12Rik | -2.29037 | 5.178364 | 38.52895 | 5.39E-10 | 5.08E-06 |
| Snord118 | -1.87849 | 0.516953 | 25.6885 | 4.01E-07 | 0.000945 |
| Dipk1b | -1.85921 | 5.720053 | 16.91818 | 3.90E-05 | 0.035016 |
| Hspa8 | -1.71342 | 8.138371 | 16.12164 | 5.94E-05 | 0.046635 |
| Pabpc4 | -1.60704 | 7.670663 | 35.52916 | 2.51E-09 | 1.58E-05 |
| AL606917.1 | -1.3759 | 2.495929 | 20.33902 | 6.49E-06 | 0.008964 |
| Ahcy | -1.36006 | 5.309481 | 34.21312 | 4.94E-09 | 2.33E-05 |
| Rab26 | -1.29743 | 6.107345 | 39.4806 | 3.31E-10 | 5.08E-06 |
| Snx5 | -1.17722 | 9.279283 | 25.18624 | 5.21E-07 | 0.00109 |
| Rfc4 | -1.10251 | 6.972421 | 24.21834 | 8.60E-07 | 0.001621 |
| Ctsl | -1.07696 | 5.785042 | 21.02822 | 4.53E-06 | 0.007106 |
| Klhl13 | -0.92559 | 5.577396 | 20.22746 | 6.88E-06 | 0.008964 |
| Dnttip1 | -0.9183 | 5.358007 | 25.72197 | 3.94E-07 | 0.000945 |
| Glul | -0.90523 | 4.723419 | 27.06426 | 1.97E-07 | 0.000742 |
| 4930444E06Rik | -0.86658 | 4.535658 | 26.48539 | 2.66E-07 | 0.000834 |
| Tubb2b | -0.86405 | 3.797749 | 20.1239 | 7.26E-06 | 0.008964 |
| P4hb | -0.79503 | 7.456829 | 21.68691 | 3.21E-06 | 0.005498 |
| Hspa5 | -0.79277 | 6.460378 | 19.69517 | 9.08E-06 | 0.009508 |
| Csrp2 | -0.69126 | 5.960674 | 19.91697 | 8.09E-06 | 0.008964 |
| Mknk2 | -0.6687 | 6.141447 | 18.08287 | 2.11E-05 | 0.020974 |
| Pdlim7 | -0.6415 | 6.236679 | 19.92807 | 8.04E-06 | 0.008964 |
| Uchl1 | -0.61761 | 6.219763 | 17.37974 | 3.06E-05 | 0.028835 |
| Traf7 | -0.57181 | 5.747142 | 16.82061 | 4.11E-05 | 0.035187 |
| Olfr1373 | 0.891491 | 2.41121 | 16.14429 | 5.87E-05 | 0.046635* |

**Supplemental Figure 3. Replicate analysis of TBP and Pol II CUT&Tag, and NET-seq data on distinct subsets of genes.**

**(A)** Average plots (top) and heatmaps arranged by decreasing Pol II occupancy (bottom) of Pol II CUT&Tag (green) and NET-seq (blue) on a 2kb window surrounding the TSS of all TATA and TATAless genes for control and IAA-treated C64 mESCs with two biological replicates each. **(B)** Top upregulated/downregulated genes extracted from edgeR analysis of RNA Pol II CUT&Tag on all genes in control and IAA-treated C64 mESCs ordered from most downregulated (top) to upregulated (bottom). Based on Figure 2C-D. **(C)** Gene browser tracks at *Rab26* (left) and *Eif4a2* (right) for CUT&Tag analyses of Pol II (green), and strand-specific reads from NET-seq data (blue) of two biological replicates of control (Con) or IAA-treated (IAA) C64 mESCs. **(D)** Average pausing index of Pol II across all genes in control (blue) and IAA treatments (red) as calculated from Pol II CUT&Tag data for control (blue) or IAA (red) C64 mESCs with two biological replicates.**(E)** Gene browser tracks for different enhancer regions for Pol II CUT&Tag analysis of two biological replicates in control control (Con) or IAA-treated C64 mESCs.

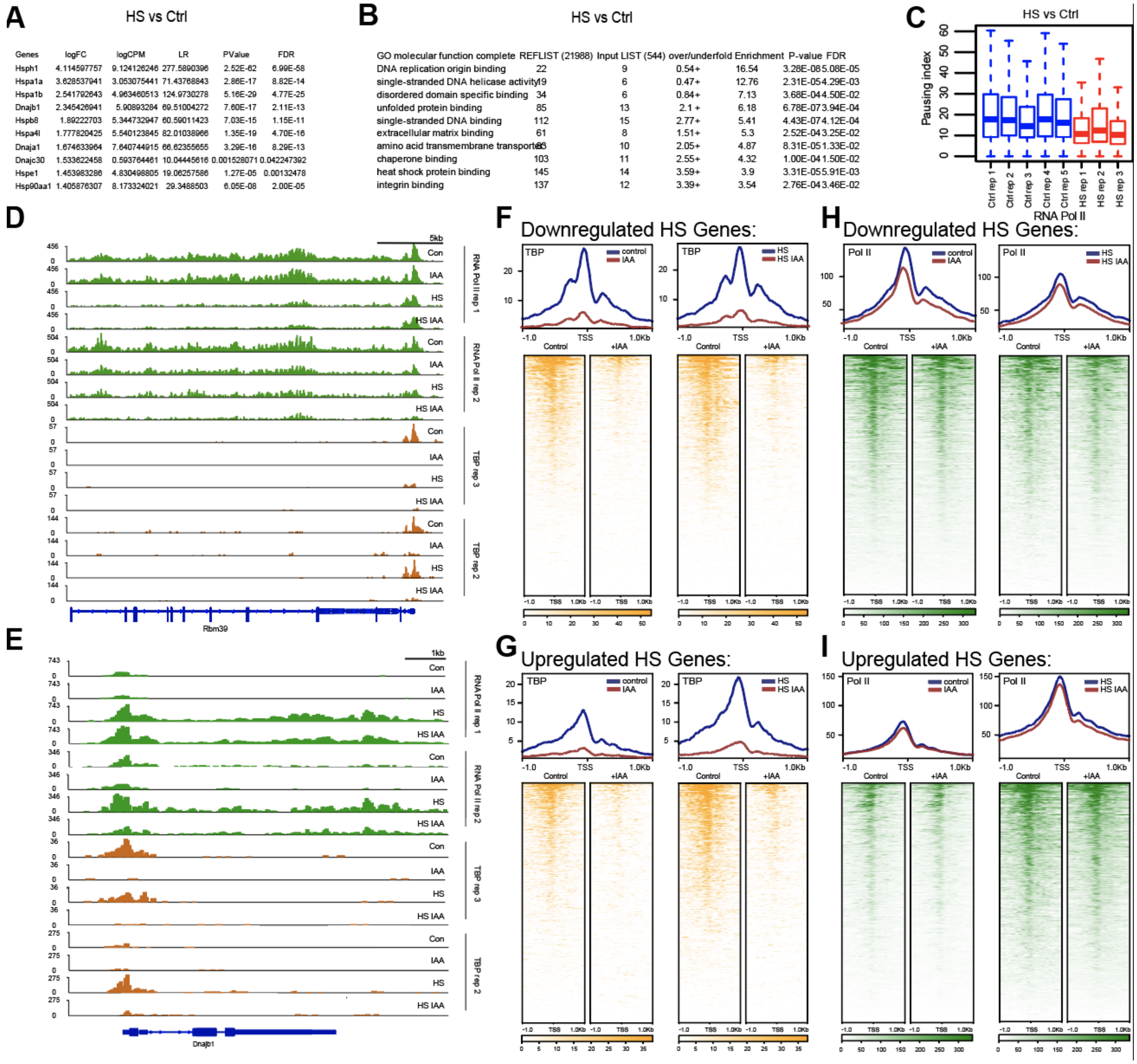

**A** HS vs Ctrl

| Genes | logFC | logCPM | LR | PValue | FDR |
|---|---|---|---|---|---|
| Hsph1 | 4.114597757 | 9.124126246 | 277.5890396 | 2.52E-62 | 6.99E-58 |
| Hspa1a | 3.628537941 | 3.053075441 | 71.43768843 | 2.86E-17 | 8.82E-14 |
| Hspa1b | 2.541792643 | 4.963460513 | 124.9730278 | 5.16E-29 | 4.77E-25 |
| Dnajb1 | 2.345426941 | 5.90893284 | 69.51004272 | 7.60E-17 | 2.11E-13 |
| Hspb8 | 1.89222703 | 5.344732947 | 60.59011423 | 7.03E-15 | 1.15E-11 |
| Hspa4l | 1.777820425 | 5.540123845 | 82.01038966 | 1.35E-19 | 4.70E-16 |
| Dnaja1 | 1.674633964 | 7.640744915 | 66.62355655 | 3.29E-16 | 8.29E-13 |
| Dnajc30 | 1.533622458 | 0.593764461 | 10.04445616 | 0.001528071 | 0.042247392 |
| Hspe1 | 1.453983286 | 4.830498805 | 19.06257586 | 1.27E-05 | 0.00132478 |
| Hsp90aa1 | 1.405876307 | 8.173324021 | 29.3488503 | 6.05E-08 | 2.00E-05 |

**B** HS vs Ctrl

| GO molecular function complete | REFLIST (21988) | Input LIST (544) | over/underfold | Enrichment | P-value | FDR |
|---|---|---|---|---|---|---|
| DNA replication origin binding | 22 | 9 | 0.54+ | 16.54 | 3.28E-08 | 5.08E-05 |
| single-stranded DNA helicase activity | 19 | 6 | 0.47+ | 12.76 | 2.31E-05 | 4.29E-03 |
| disordered domain specific binding | 34 | 6 | 0.84+ | 7.13 | 3.68E-04 | 4.50E-02 |
| unfolded protein binding | 85 | 13 | 2.1 + | 6.18 | 6.78E-07 | 3.94E-04 |
| single-stranded DNA binding | 112 | 15 | 2.77+ | 5.41 | 4.43E-07 | 4.12E-04 |
| extracellular matrix binding | 61 | 8 | 1.51+ | 5.3 | 2.52E-04 | 3.25E-02 |
| amino acid transmembrane transporter | [illegible] | 10 | 2.05+ | 4.87 | 8.31E-05 | 1.33E-02 |
| chaperone binding | 103 | 11 | 2.55+ | 4.32 | 1.00E-04 | 1.50E-02 |
| heat shock protein binding | 145 | 14 | 3.59+ | 3.9 | 3.31E-05 | 5.91E-03 |
| integrin binding | 137 | 12 | 3.39+ | 3.54 | 2.76E-04 | 3.46E-02 |



### Supplemental Figure 4. TBP is not required for induction of heat shock genes.

**(A)** Raw values of a few top upregulated HS genes extracted from edgeR analysis of RNA Pol II CUT&Tag on all genes in control and HS-treated C64 mESCs ordered from most upregulated to least. Based on Figure 3C. **(B)** Gene ontology of the top upregulated genes extracted from edgeR analysis of RNA Pol II CUT&Tag on all genes in control and HS-treated C64 mESCs.**(C)** Average pausing index of Pol II across top upregulated HS genes extracted from edgeR analysis in control (blue) and HS samples (red) as calculated from Pol II CUT&Tag data for individual replicates (5 Control, 3 Heat shock) in C64 mESCs. **(D-E)** Gene browser tracks for *Rbm39* (top) and *Dnajb1* (bottom) for CUT&Tag analyses of Pol II (green) and TBP (orange) in control, IAA-treated, HS and HS IAA treated C64 mESCs for two biological replicate. **(F-G)** Average plots (top) and heatmaps arranged by decreasing TBP occupancy (bottom) of TBP CUT&Tag in a 2kb window surrounding the TSS of the top downregulated (top) or top upregulated (bottom) HS genes identified from edgeR for control, IAA, HS, and HS IAA-treated C64 mESCs. **(H-I)** Average plots (top) and heatmaps arranged by decreasing Pol II occupancy (bottom) of Pol II CUT&Tag in a 2kb window surrounding the TSS of the top downregulated (top) or top upregulated (bottom) HS genes identified from edgeR for control, IAA, HS, and HS IAA-treated C64 mESCs.

A

RA vs Ctrl

| Genes | logFC | logCPM | LR | PValue | FDR |
|---|---|---|---|---|---|
| Tdpoz4 | 6.453303 | -1.89779 | 19.45969 | 1.03E-05 | 0.002479 |
| Stra8 | 3.265719 | 6.30867 | 111.1632 | 5.45E-26 | 5.04E-22 |
| Nrip1 | 2.390513 | 6.625802 | 129.8103 | 4.51E-30 | 1.25E-25 |
| Pbx1 | 2.298765 | 7.485449 | 124.0998 | 8.01E-29 | 1.11E-24 |
| Rbp1 | 1.98741 | 3.961606 | 67.94815 | 1.68E-16 | 6.65E-13 |
| Cdx1 | 1.39898 | 4.216878 | 29.27348 | 6.28E-08 | 4.15E-05 |
| Ski | 1.354878 | 8.534942 | 41.43548 | 1.22E-10 | 1.69E-07 |
| Zfhx3 | 1.235738 | 9.01187 | 28.19229 | 1.10E-07 | 5.98E-05 |
| Foxp1 | 1.205978 | 8.485204 | 22.72876 | 1.87E-06 | 0.000631 |

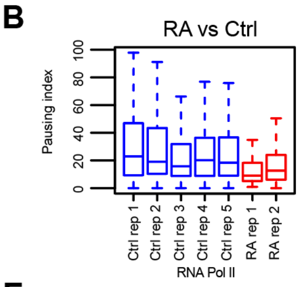


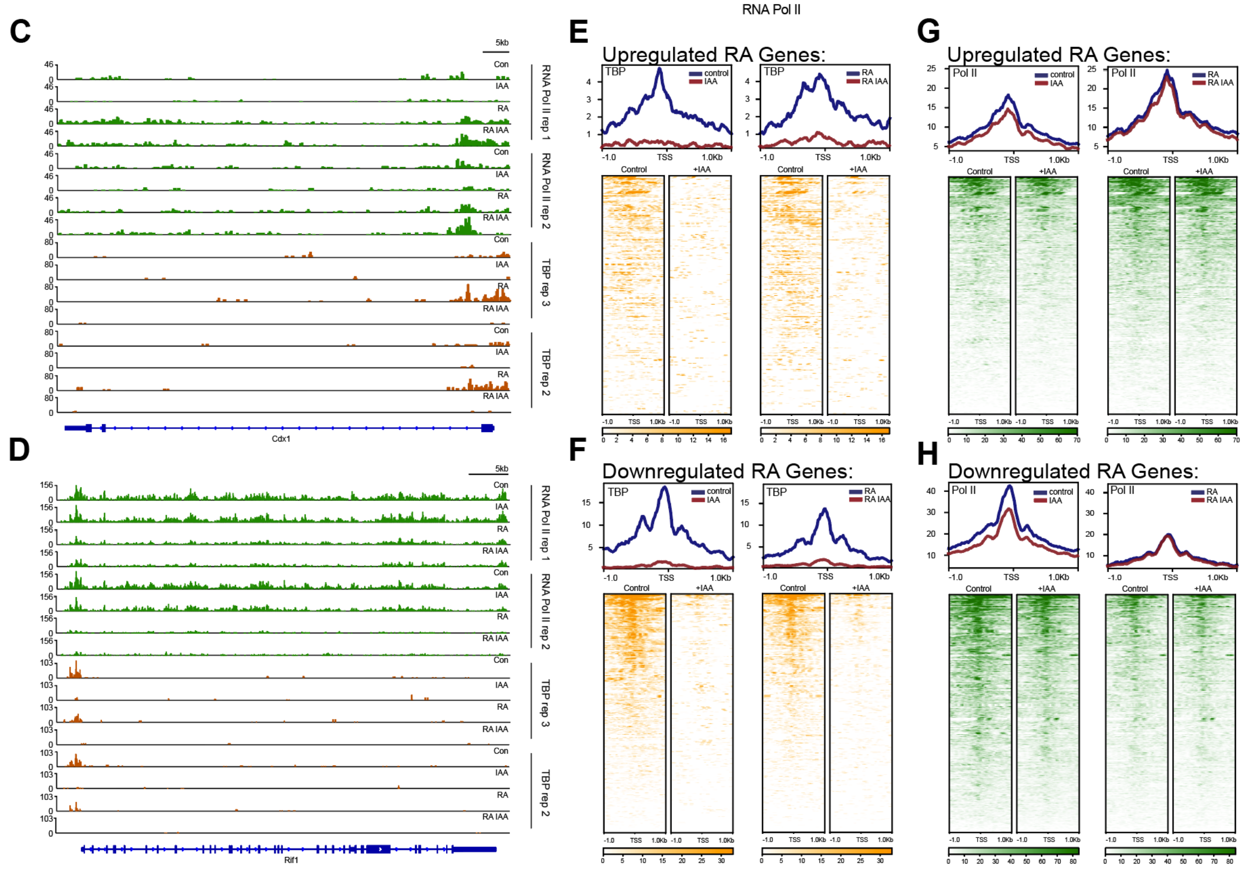


### Supplemental Figure 5. TBP is not required for induction of RA-differentiation genes.

**(A)** Raw values of a few top upregulated RA genes extracted from edgeR analysis of RNA Pol II CUT&Tag on all genes in control and RA-treated C64 mESCs ordered from most upregulated to least. **(B)** Average pausing index of Pol II across top upregulated RA genes extracted from edgeR analysis in control (blue) and RA samples (red) as calculated from Pol II CUT&Tag data for individual replicates (5 control, 2 RA) in C64 mESCs.**(C-D)** Gene browser tracks for *Cdx1* (top) and *Rif1* (bottom) for CUT&Tag analyses of Pol II (green) and TBP (orange) in control, IAA-treated, RA and RA IAA treated C64 mESCs for two biological replicate. **(E-F)** Average plots (top) and heatmaps arranged by decreasing TBP occupancy (bottom) of TBP CUT&Tag in a 2kb window surrounding the TSS of the top upregulated (top) or top downregulated (bottom) RA genes identified from edgeR for control, IAA, RA and RA IAA-treated C64 mESCs. **(H-I)** Average plots (top) and heatmaps arranged by decreasing Pol II occupancy (bottom) of Pol II CUT&Tag in a 2kb window surrounding the TSS of the top upregulated (top) or top downregulated (bottom) RA genes identified from edgeR for control, IAA, RA and RA IAA-treated C64 mESCs..

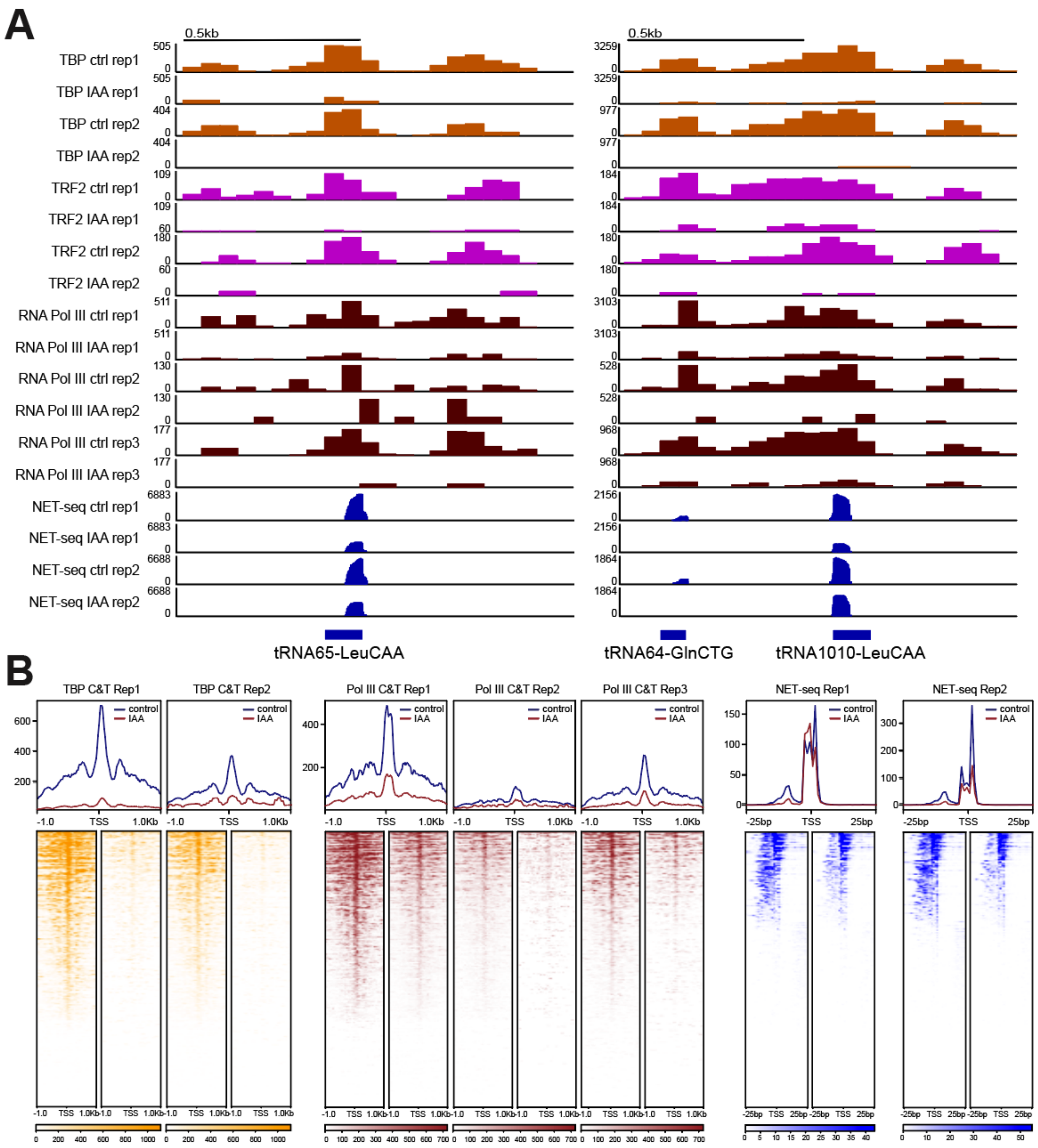


**Supplemental Figure 6. Replicate analysis of TBP and Pol III CUT&Tag on tRNA genes.**

**(A)** Gene browser tracks at tRNA65-LeuCAA (left) and tRNA64-GlnCTG (right) for CUT&Tag analyses of TBP (orange), TRF2 (magenta), Pol III (maroon), and strand-specific reads from NET-seq data (blue) in two biological replicates of control or IAA-treated C64 mESCs. **(B)** Average plots (top) and heatmaps arranged by decreasing TBP occupancy (bottom) of TBP CUT&Tag (left), Pol III CUT&Tag (middle), and NET-seq (right) in a 2kb window (CUT&Tag) or 50 bp window (NET-seq) surrounding the TSS of all tRNAs in two biological replicates of control and IAA-treated C64 mESCs.

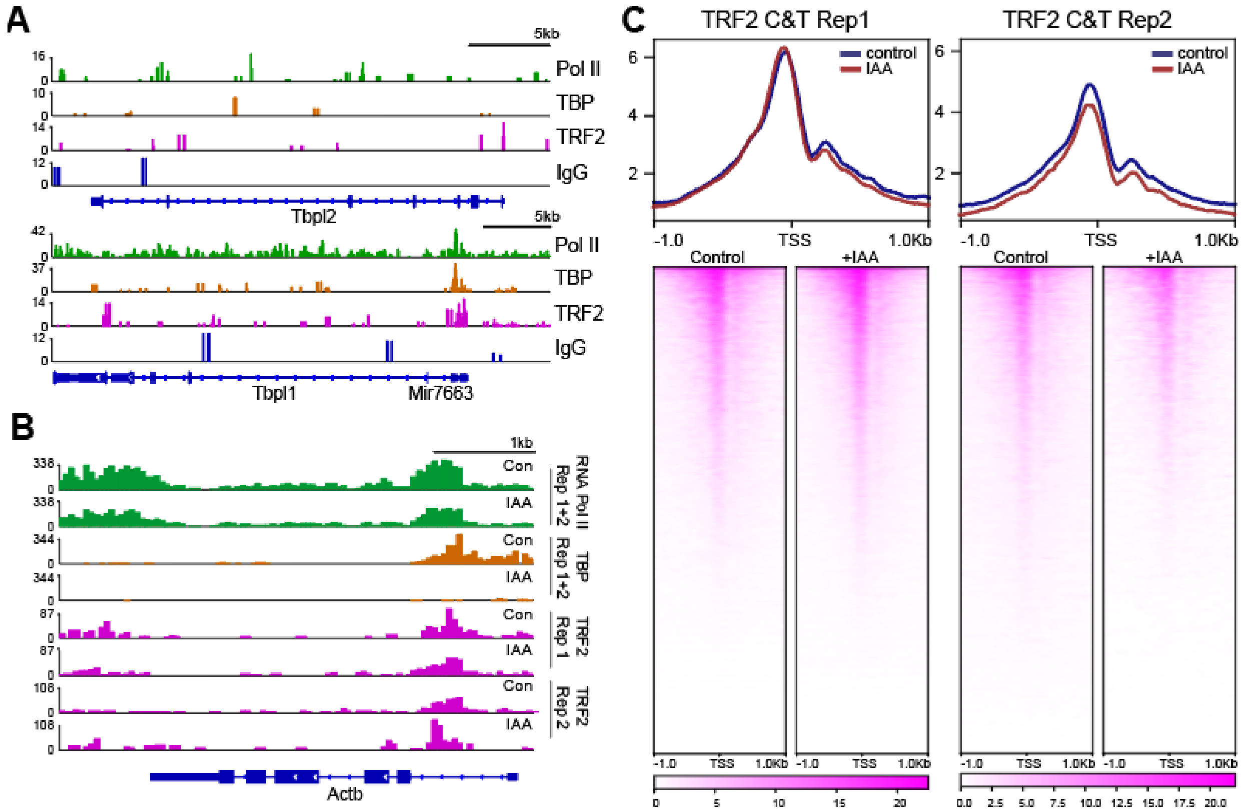


**Supplemental Figure 7. Replicate analysis of TRF2 CUT&Tag data.**

**(A)** Gene browser tracks for *Tbpl2* (top) and *Tbpl1* (bottom) for CUT&Tag analyses of Pol II (green), TBP (orange), TRF2 (magenta), and IgG (blue) in C64 mESCs. **(B)** Gene browser tracks for *Actb* for CUT&Tag analyses of Pol II (green), TBP (orange) and TRF2 (magenta) in control and IAA-treated C64 mESCs with two biological TRF2 replicas. **(C)** Average plots (top) and heatmaps arranged by decreasing RNA Pol II occupancy (bottom) of TRF2 CUT&Tag in a 2kb window CUT&Tag surrounding the TSS of all genes for two biological replicates of control and IAA-treated C64 mESCs.

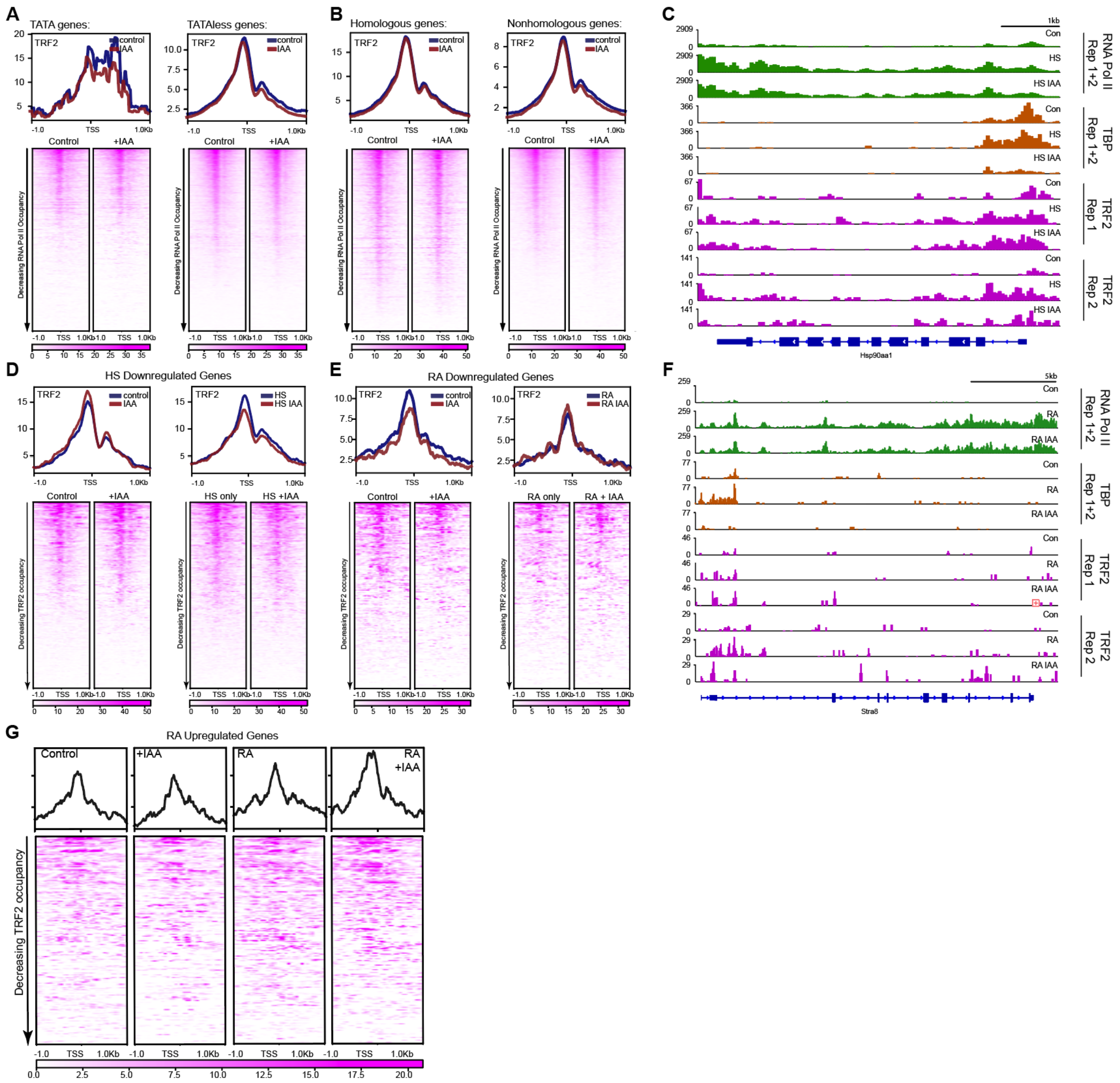


**Supplemental Figure 8. TRF2 binding on distinct subsets of genes and on induced genes.**

**(A)** Average plots (top) and heatmaps arranged by decreasing RNA Pol II occupancy (bottom) of TRF2 CUT&Tag in a 2kb window CUT&Tag surrounding the TSS of TATA genes (left) or TATAless genes (right) for control and IAA-treated C64 mESCs. **(B)** Average plots (top) and heatmaps arranged by decreasing RNA Pol II occupancy (bottom) of TRF2 CUT&Tag in a 2kb window CUT&Tag surrounding the TSS of Homologous genes (left) or Non-Homologous genes (right) for control and IAA-treated C64 mESCs. **(C, F)** Gene browser tracks for *Hsp90aa1* **(C)** and *Stra8* **(F)** for CUT&Tag analyses of Pol II (green) TBP (orange) and TRF2 (magenta) in control, HS and HS IAA-treated (top), or control, RA and RA IAA-treated (bottom) C64 mESCs with two biological TRF2 replicas. **(D)** Average plots (top) and heatmaps arranged by decreasing TRF2 occupancy (bottom) of TRF2 CUT&Tag in a 2kb window surrounding the TSS of the top upregulated HS genes identified from edgeR for control, IAA, HS and HS IAA-treated C64 mESCs. **(E)** Average plots (top) and heatmaps arranged by decreasing TRF2 occupancy (bottom) of TRF2 CUT&Tag in a 2kb window surrounding the TSS of the top upregulated RA genes identified from edgeR for control, IAA, RA and RA IAA-treated C64 mESCs. **(G)** Average plots (top) and heatmaps arranged by decreasing TRF2 occupancy (bottom) of TRF2 CUT&Tag in control, IAA-treated (+IAA), retinoic acid treated (RA), and RA and IAA-treated (RA + IAA) C64 mESCs in a 2kb window surrounding the TSS of all genes upregulated RA treatment.
